# Mapping the invisible: Incidental spatial encoding through haptic exploration

**DOI:** 10.64898/2026.11.24.768822

**Authors:** Maria Evangelia Vlachou, Sara Kasim, Laurence Mouchnino, Jean Blouin

**Author notes:** **Corresponding author:** Jean Blouin, Institut des Sciences du Mouvement Etienne Jules Marey (ISM), Aix-Marseille Université, 163 Avenue de Luminy, 13288 Marseille Cedex 09, France.

## Abstract

Active touch provides rich spatial information about object shape. Whether this haptic information can be retained without an explicit memorisation goal remains unclear. Participants explored an unfamiliar two-dimensional raised-dot contour under different sensory conditions and memorisation instructions. Two haptic groups traced the contour hidden from view, either without advance knowledge of the memory tasks (Haptic-Naïve) or with explicit memorisation instructions (Haptic-Informed). Reference groups traced the visible contour (Visuo-Haptic-Naïve) or viewed it without tracing (Visual-Naïve), both without memorisation instructions. Retention was assessed through manual shape reproduction and visual recognition, with a Random-Control group drawing arbitrary shapes as a reproduction baseline. Reproductions in all exploration groups were more similar to the target than arbitrary drawings, supporting incidental spatial encoding in the Haptic-Naïve group. Memorisation instructions produced no detectable reproduction advantage but altered exploratory behaviour. Visual feedback improved online tracing without a detectable reproduction advantage over haptic-only exploration. Visual-Naïve participants reproduced the shape more accurately than Haptic-Naïve participants. Recognition scores were higher following memorisation instructions or exploration with visual information, although Haptic-Naïve participants selected alternatives relatively similar to the target. Source-space EEG analyses in the Haptic-Naïve group revealed widespread beta-power reductions across bilateral somatosensory, parietal, and occipital regions. Greater reductions in the left lateral occipital complex were associated with more accurate subsequent reproduction. Together, these findings indicate that active touch can support incidental spatial encoding and is associated with changes in cortical activity. The accessibility and use of the retained information are shaped by the goals and sensory context of exploration.

## Introduction

Successful navigation and object interaction rely on internal spatial representations that can be maintained in working memory and used to guide goal-directed behaviour. Vision is often the dominant source of such information, yet it is not indispensable. When visual information is limited or absent, touch can also extract spatial structure (Klatzky et al., 1985, 1993), albeit with reduced accuracy and slower processing compared to vision (Loomis et al., 1991).

The ability of touch to support spatial perception has been attributed to its dual nature, integrating both allocentric and egocentric information. Cutaneous feedback conveys fine-grained information about contacted surfaces, including their edges, contours, and textures (i.e., allocentric information; Johnson, 2001), while also providing cues about hand displacement and movement (i.e., egocentric information; Moscatelli et al., 2019; Vlachou et al., 2025). Kinaesthetic feedback from muscles, tendons, and joints provides additional information about hand position and movement. Combining these signals helps relate successive contacts to one another, supporting the perception of spatial properties such as curvature, shape, and orientation during active exploration (Gibson, 1966; Loomis & Lederman, 1986). The relative contributions of cutaneous and kinaesthetic information depend on stimulus properties and task demands (Symmons et al., 2004). For example, cutaneous input alone can support the identification of small tactile figures, whereas movement-related proprioceptive information becomes more important for accurately identifying larger figures (Loo et al., 1983).

A well-established paradigm for studying haptic spatial perception involves exploring two-dimensional raised-line drawings depicting familiar objects (Lebaz et al., 2012; Lederman et al., 1990; Loomis et al., 1991; Wijntjes et al., 2008) or basic geometric forms (Heimler et al., 2021; Marlair et al., 2021). When the explored configurations are familiar, their perception can be supported by previously acquired knowledge about their identity and structure. Such knowledge may contribute to haptic perception through mental imagery. A distinction is commonly made between object imagery, which emphasises an object’s pictorial appearance, and spatial imagery, which emphasises spatial relationships among objects or their component parts. Both can involve visual representations (Blajenkova et al., 2006). Neuroimaging findings suggest that the relationship between haptic perception and these forms of imagery varies with object familiarity. Neural activity during haptic perception of familiar objects showed more extensive overlap and stronger correlations with activity during visual object imagery than did activity during haptic perception of unfamiliar objects (Lacey et al., 2010). Effective-connectivity analyses similarly indicated a closer correspondence between the networks underlying visual object imagery and haptic perception of familiar objects (Deshpande et al., 2010). By comparison, the network underlying spatial imagery shared more effective-connectivity pathways with haptic perception of unfamiliar than familiar objects (Lacey et al., 2014). Together, these findings suggest that processing unfamiliar shapes, which do not readily evoke a known object identity, may depend more strongly on integrating spatial information acquired during exploration. Unfamiliar shapes may therefore be particularly useful for examining spatial encoding through touch while reducing the contribution of prior object knowledge.

Despite their potential for examining spatial encoding, relatively few studies have used unfamiliar shapes, and these studies have generally informed participants in advance that their memory for the explored patterns would be tested (Bailes & Lambert, 1986; Behrmann & Ewell, 2003; Picard et al., 2010). In these tasks, participants reported using spatial, visuo-spatial, verbal, and kinaesthetic encoding strategies (Picard et al., 2010; Picard & Monnier, 2009). However, retention does not necessarily require an explicit memorisation goal. In vision, detailed information about previously viewed objects can be retained incidentally following visual search (Castelhano & Henderson, 2005), although subsequent reproduction can be less accurate when participants do not anticipate the memory task (Cárdenas-Miller et al., 2025). Similar evidence comes from haptic exploration of everyday objects. Participants who explored objects while blindfolded, without prior memorisation instructions, subsequently recognised them through either touch or vision in unexpected memory tests one week later (Hutmacher & Kuhbandner, 2018). These findings demonstrate incidental retention of everyday objects, but leave open whether haptic exploration of unfamiliar shapes that do not readily evoke a known object identity can similarly support the encoding of their spatial configuration. Addressing this question would help clarify the extent to which haptic spatial encoding can occur spontaneously and the limits of this capacity.

In the present study, we investigated whether haptic exploration without prior memorisation instructions supports spatial encoding of an unfamiliar two-dimensional shape defined by a raised-dot contour. Retention was assessed through both subsequent manual reproduction and visual recognition. Participants in the Haptic-Naïve (H-N) group traced the contour while it remained hidden from view, without advance knowledge of the memory tasks. The Haptic-Informed (H-I) group performed the same exploration task with explicit memorisation instructions. Comparing these groups allowed us to assess the capacity of tactile exploration to support incidental encoding of spatial information in the absence of an explicit memorisation goal. Three additional groups served as reference conditions. The Visuo-Haptic-Naïve (VH-N) group traced the visible contour, whereas the Visual-Naïve (V-N) group viewed it without tracing. Neither group received prior memorisation instructions, allowing us to assess incidental retention following visual exploration, either with the sensorimotor activity associated with tracing (VH-N) or without it (V-N). The Random-Control (RC) group drew arbitrary shapes without prior exposure to the target, providing a baseline against which to evaluate reproduction following exploration.

We hypothesised that, if haptic exploration supports incidental spatial encoding, reproductions in the H-N group would be more similar to the target than arbitrary drawings produced by the RC group. We further hypothesised that repeated tracing would support sufficient incidental encoding for subsequent reproduction and recognition, with no additional benefit from explicit memorisation instructions. We therefore predicted comparable performance in the H-N and H-I groups. Given the prominent role of vision in spatial processing, we were also particularly interested in comparing retention following haptic-only exploration with that observed when visual information about the shape was available, either alone or combined with haptic exploration.

Finally, we recorded EEG during tracing in the H-N group to characterise cortical activity during haptic exploration. Neuroimaging studies implicate somatosensory and parietal regions, together with the lateral occipital complex (LOC), in haptic processing of unfamiliar shapes (Deshpande et al., 2010; Lacey & Sathian, 2014). We focused on beta-band activity given its stronger modulation during touch-guided than visually guided tracing (Lee & Kukke, 2017) and its involvement in tactile working memory (Li Hegner et al., 2007). Using source-space spectral analyses, we quantified task-related beta-power changes and examined their relationships with the subsequent memory tasks.

## Materials and Methods

### Participants

A total of 121 healthy adults (mean age ± SD = 24.6 ± 3.3 years) participated in the study. All participants were right-handed, as assessed by the Edinburgh Handedness Inventory (Oldfield, 1971) (mean laterality score ± SD = 80.7 ± 23.2), and reported no visual, motor, or neurological impairments. Participants were randomly assigned to one of five experimental groups: Haptic-Naïve (H-N) (16 men, 21 women), Haptic-Informed (H-I) (7 men, 14 women), Visuo-Haptic-Naïve (VH-N) (4 men, 15 women), Visual-Naïve (V-N) (4 men, 16 women), and Random-Control (RC) (5 men, 19 women). The H-N group included a larger number of participants than the other groups because it also served as the EEG cohort, allowing sufficient statistical power for the neurophysiological analyses. Experimental sessions lasted ∼60 min for the H-N group and ∼20 min for the other groups, which did not require EEG preparation and recording. Participants in the H-N group received €20 upon completion of the experiment. The study was conducted in accordance with the principles of the Declaration of Helsinki (2013 revision) and was approved by the CERSTAPS ethics committee (approval no. IRB00012476-2021-09-12-140). Written informed consent was obtained from all participants prior to participation.

### Experimental set-up and stimulus

Participants were seated comfortably at a table in a dimly lit room. Positioned in front of them was a force plate (HE6×6, AMTI, Watertown, MA, USA; supplied by A-Tech Instruments Ltd, Toronto, ON, Canada; 150 × 150 mm) supporting a black biopolymer thermoplastic surface (polylactic acid, PLA; 195 × 195 × 2 mm). On this surface, the white contour of an unfamiliar closed irregular polygon, hereafter referred to as the Reference shape, was 3D-printed (Sigma D25 printer, BCN3D Technologies, Gavà, Barcelona, Spain) (Fig. 1A). The Reference shape had a total perimeter of 652 mm and consisted of 17 linear segments, each 5 mm wide. Its contour was defined by three parallel rows of raised Braille-like dots arranged in a hexagonal 7-dot configuration (Fig. 1B). Each dot measured 1.5 mm in diameter and 0.6 mm in height, with an inter-dot spacing of 2.5 mm. This texture was designed to provide consistent tactile stimulation of cutaneous mechanoreceptors involved in spatial discrimination (K. O. Johnson & Lamb, 1981). The surrounding surface was smooth and featureless, providing no additional spatial tactile cues outside the shape.

**Figure 1.**
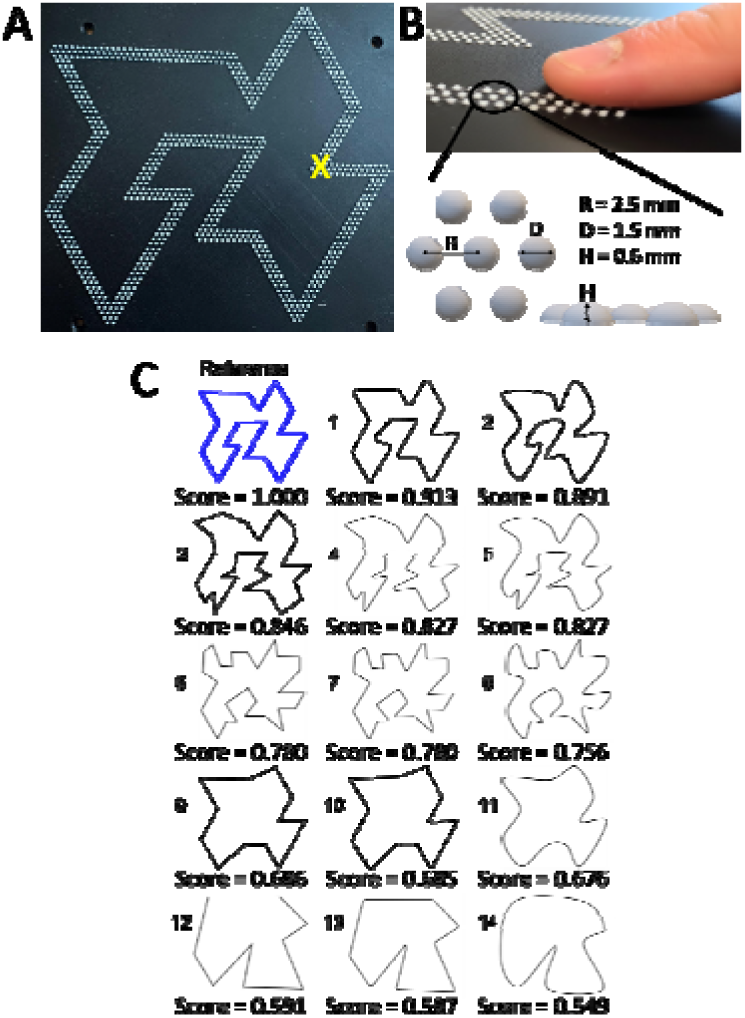
Experimental setup and stimuli. **A** Exploration surface with the contour of the Reference shape 3D-printed as a raised, Braille-like dot pattern. The X symbol, shown for illustrative purposes only, indicates the starting point of the exploration phase and was not present on the actual surface. **B** Close-up of the fingertip interacting with the Braille-like dot contour and schematic of the dot configuration. The contour was formed by repeating clusters of seven white dots arranged in a hexagonal pattern (dot diameter: 1.5 mm; height: 0.6 mm; inter-dot spacing: 2.5 mm), printed on a smooth, featureless black surface. **C** Reference shape (top left) and the set of candidate shapes from which participants selected up to three during the recognition task. For illustration, the candidate shapes are displayed here in descending order of similarity to the Reference, as quantified by the Geometric–Feature Similarity (GFS) algorithm. The candidate shapes were numbered 1-14 according to this similarity ranking. During the experiment, the shapes were presented on a sheet of paper (21 × 29.7 cm) in randomised order, without numbering or similarity scores.

### Experimental groups

Participants were randomly assigned to one of five experimental groups (Fig. 2A) that differed in sensory modality during exploration and in whether they received prior information about the subsequent tasks (Fig. 2B). The Haptic-Naïve (H-N) group traced the Reference shape with the index fingertip, while a black shield prevented vision of both the hand and the shape. Participants kept their eyes open and were not informed that they would later be asked to both reproduce the shape and select it from a set of candidate shapes. Their only instruction was to follow the tactile contour. The Haptic-Informed (H-I) group performed the same haptic tracing task but received explicit instructions to memorise the shape before exploration. This group served as a reference for assessing the extent of incidental spatial encoding in the H-N group following exploration without an explicit memorisation goal.

**Figure 2.**
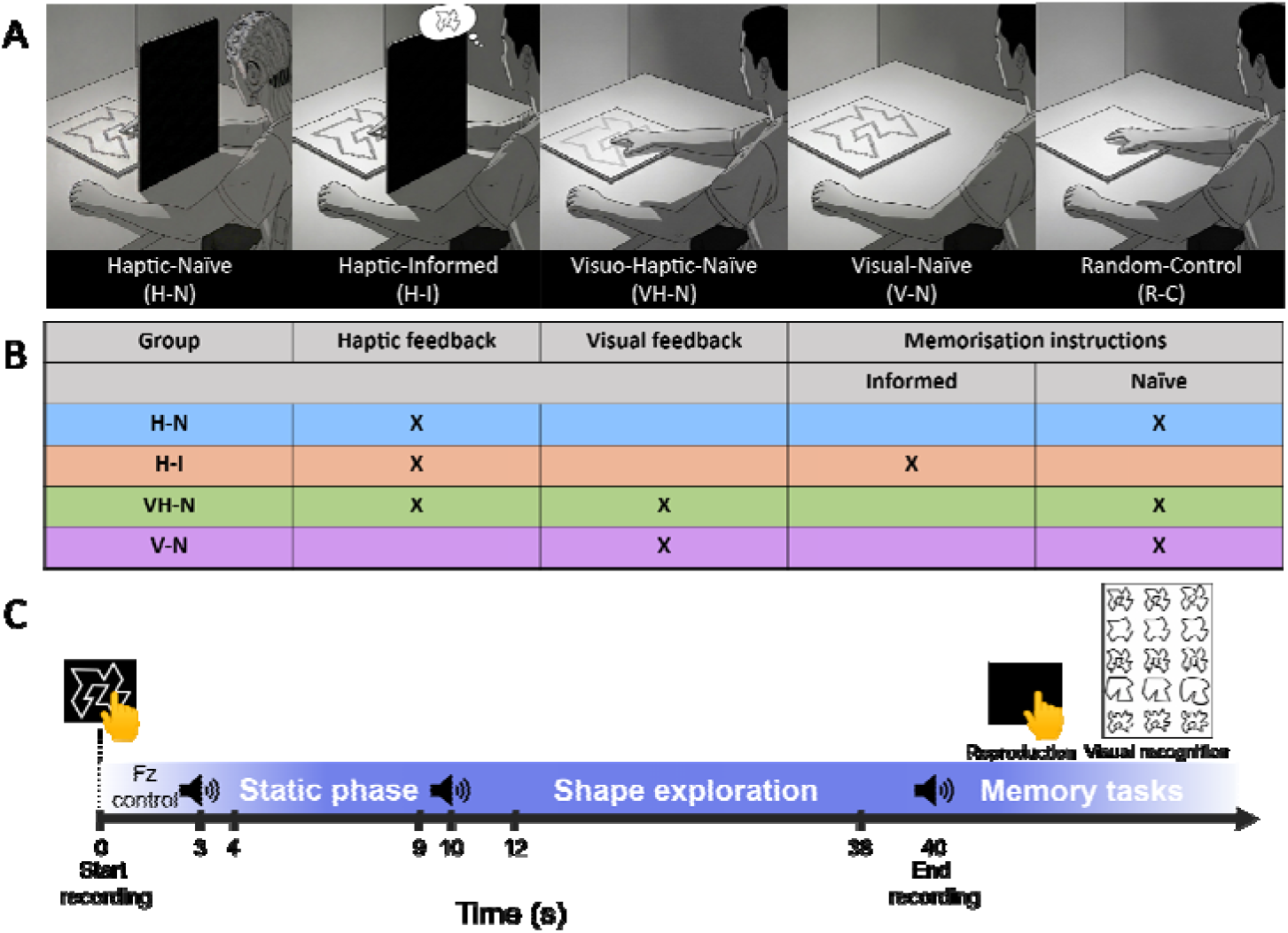
Experimental design and trial structure. **A** Schematic overview of the five experimental groups. **B** Summary table of the sensory information available during exploration (haptic and/or visual) and whether participants received prior memorisation instructions. The RC group is not represented in the table because participants in this group neither explored nor had prior exposure to the Reference shape; instead, they drew an arbitrary closed shape. **C** Temporal organisation of the exploration phase and subsequent memory tasks. Exploration consisted of 20 consecutive trials. In the haptic groups (H-N, H-I and VH-N), each trial comprised a 3-s period of normal force stabilisation, a static baseline (7 s for H-N and 3 s for H-I and VH-N), and a 30-s tracing period. Participants resumed tracing from the endpoint of the preceding trial, allowing sequential exploration of the entire contour. Participants in the V-N group viewed the Reference shape continuously throughout each 36-s trial without tracing. For the H-N group, beta-band spectral analyses used the 12–38 s interval during tracing and the 4–9 s interval during the static baseline, with both intervals defined relative to trial onset. After exploration, participants completed two memory tasks involving manual reproduction of the shape with the index finger on a smooth black surface and visual recognition among a set of candidate shapes.

With the black shield removed, the Visuo-Haptic-Naïve (VH-N) group traced the shape with simultaneous visual and tactile feedback but, like the H-N group, was unaware of the subsequent tasks. The Visual-Naïve (V-N) group viewed the shape without tracing and likewise received no information about the subsequent tasks. To ensure equivalent exposure time across groups, the Reference shape was occluded before and between trials in both the VH-N and V-N groups. Finally, unlike participants in the other groups, those in the Random-Control (RC) group did not explore the Reference shape. Instead, their only task was to draw an arbitrary closed shape with full vision, without prior acquisition of shape-specific spatial information. This provided a baseline against which the reproduction performance of the other groups could be evaluated.

### Procedure and task phases

Before the experiment, participants washed and thoroughly dried their hands. For all groups except the RC group, the experimental protocol consisted of three consecutive phases: exploration, reproduction, and recognition.

During the exploration phase, participants explored the Reference shape according to their assigned group over 20 trials. In the haptic groups, the participant’s index fingertip was positioned at the designated starting point before the first trial (X mark in Fig. 1A). In the H-N and H-I groups, in which a black shield prevented vision of both the shape and the hand, the experimenter positioned the participant’s fingertip. Trials in the haptic groups (H-N, H-I and VH-N) began with a 3-s force-adjustment period, during which participants stabilised the normal force applied to the surface within a range of 0.2–0.4 N, consistent with natural tactile exploration (Smith et al., 2002). Force levels were monitored in real time on the experimenter’s display, and corrective feedback was provided when necessary. Following the normal force stabilisation period, an auditory cue (“beep”) marked the onset of a static baseline period, during which participants remained motionless while maintaining the normal force against the surface.

The baseline period lasted 7 s for the H-N group, providing a reliable baseline for the subsequent EEG analyses, and 3 s for the H-I and VH-N groups, in which EEG was not recorded. A second beep marked the start of the 30-s exploration period, and a third beep signalled the end of the trial (Fig. 2C). Trials followed one another without an additional pause, with each new trial beginning with the 3-s force-adjustment period. During the exploration period, participants in the H-N, H-I, and VH-N groups traced the contour of the 3D-printed shape. In contrast, to match the visual exposure of the VH-N group, participants in the V-N group viewed the Reference shape continuously throughout the entire 36-s trial (baseline + exploration period), without tracing.

In the haptic groups, participants were encouraged to trace the contour slowly (∼10 mm/s), although no strict speed constraint was imposed. This target speed was selected for the H-N group to minimise potential contamination of the EEG signal by muscle and ocular artefacts while allowing the shape to be explored progressively across trials. At this speed, the full contour could not be completed within a single trial. Each trial therefore resumed from the point at which tracing had ended in the previous trial. This segmented protocol provided a sufficient number of baseline– movement epochs for trial-averaged EEG analyses (see below).

After the exploration phase, participants in all four exploration groups received detailed instructions for the reproduction task. Only participants in the H-I group had been informed in advance that this task would follow. Participants then reproduced the Reference shape from memory by tracing it once with their right index finger on a smooth, textureless black surface. Participants could see their hand and its movements throughout reproduction. This was followed by a visual recognition task in which participants were presented with a printed sheet containing 15 shapes arranged in a randomised order. The set included the Reference shape and comprised five base configurations, each presented in three variants with increasing contour curvature. Participants were instructed to identify the previously explored Reference shape and could select up to three candidates if uncertain. The number of selections was interpreted as an index of recognition confidence, with one choice indicating the highest confidence and three choices the lowest.

### Kinematic and force analyses

#### Data acquisition

Finger movements were recorded using a GoPro HERO Session camera (Full HD, 1920 × 1080 pixels; 60 fps) mounted 290 mm above the exploration surface and aligned parallel to it. A small cross drawn on the participant’s fingernail served as the tracking marker for motion analysis using Kinovea (Charmant, 2024). Spatial calibration was verified by comparing the perimeter of the polygon reconstructed from the recordings with its known physical dimensions, ensuring accurate spatial scaling.

Reaction forces and moments exerted by the index finger were recorded at 256 Hz using an AMTI HE6×6 force plate mounted beneath the 2-mm-thick exploration surface. The system provided 12-bit measurement resolution, with measurement ranges of 2.2 N for Fx and Fy and 4.5 N for Fz, allowing the low-amplitude forces generated during fingertip–surface interaction to be captured. Force signals were digitised using a Keithley 12-bit A/D converter (AD-win pro, Keithley Instruments, Cleveland, OH, USA) and analysed offline in MATLAB 7.0 (The MathWorks, Natick, MA, USA).

#### Data processing and extracted measures

For each of the 20 trials performed by the haptic groups (H-N, H-I, and VH-N), kinematic and force signals were analysed from movement onset following the auditory “start tracing” cue to movement offset following the “stop tracing” cue. Five measures were extracted to characterise tracing behaviour and analysed separately from subsequent memory performance: the distance-to-segment index, number of shape traversals, tracing speed, normal force (Fz), and jerk index, as described below.

#### Kinematic measures

Tracing accuracy during the exploration phase was quantified using the distance-to-segment index, a commonly used measure of tracing performance (Lebar et al., 2015, 2017; Mathieu et al., 2023; Miall & Cole, 2007; Vlachou et al., 2025). The index was defined as the ratio between the total path length traced by the participant’s finger and the true length of the corresponding contour segment. A value of 1 indicates that the traced path matches the target segment length and is consistent with accurate tracing, whereas values greater than 1 reflect larger trajectories and reduced tracing accuracy. To compare the extent of contour exploration across the haptic groups, we estimated the number of equivalent contour traversals by dividing the cumulative contour length explored across all 20 trials by the perimeter of the Reference shape (652 mm).

Tracing speed was computed from x–y positional coordinates. Raw coordinates were pre-processed using a fourth-order, zero-phase Butterworth low-pass filter (6 Hz cutoff). Instantaneous speed was then calculated as the Euclidean displacement between successive samples divided by the fixed sampling interval 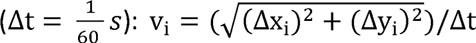. Mean tracing speed was obtained by averaging the instantaneous speed across the active movement window.

#### Force measures

Forces exerted on the surface were analysed along all three axes. Normal force (Fz) was calculated as the mean vertical force during the exploratory phase of each trial. Tangential forces in the horizontal (XY) plane were analysed using a jerk index derived from temporal changes in their resultant magnitude, providing a measure of movement smoothness:

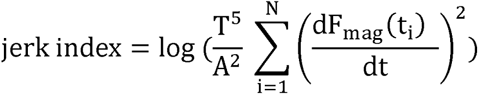

where 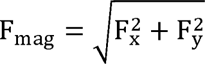 represents the resultant horizontal force magnitude, T is the trial duration, A is the force amplitude (maximum - minimum F_mag_ within each trial), and dF_mag_⁄dt is its temporal derivative. This metric captures rapid, transient changes in force production associated with corrective adjustments during shape exploration while normalising for both trial duration and force amplitude. Higher jerk index values indicate more intermittent and less stable exploratory force patterns, characterised by frequent corrective adjustments during tracing of the dotted contour. In contrast, lower jerk values indicate smoother and more continuous force modulation. Jerk-based measures have been widely used to characterise motor performance in both clinical and experimental contexts (Balasubramanian et al., 2015; Goldvasser et al., 2001; Maura et al., 2023; Vlachou et al., 2025; Vlachou, Lafaverges, et al., 2026).

### Shape similarity analysis

To quantify the similarity between the shapes reproduced by participants in each group and the Reference shape, we used the Geometric–Feature Similarity (GFS) algorithm proposed by Vlachou *et al*. (2026b). The GFS algorithm was developed to address limitations of existing shape similarity measures based on either classical geometric methods (e.g., Procrustes analysis and Chamfer distance) or on convolutional neural network (CNN)-inspired feature-based representations. It combines geometric measures, such as Chamfer distance and perimeter similarity, with convolutional feature-based descriptors capturing local structural characteristics of the shape into a single similarity measure. This combined approach more accurately reproduced the visually defined perceptual ranking of the Reference and candidate shapes than either the geometric or feature-based approaches alone, while avoiding the need for network training or large labelled datasets. A detailed description of the GFS algorithm and its comparison with other shape similarity methods is provided by Vlachou *et al*. (2026*b*).

For the reproduction task, the global GFS score was computed by comparing each participant’s reproduced shape with the Reference shape. This score served as the measure of reproduction performance, with higher values indicating greater similarity to the Reference and a score of 1 indicating perfect reproduction.

For the recognition task, each of the 15 candidate shapes was assigned a GFS score quantifying its similarity to the Reference shape (Fig. 1C). Recognition performance was quantified as the mean GFS score of the one to three shapes selected by each participant, with scores closer to one indicating a closer match to the Reference shape (GFS = 1).

### Electroencephalography (EEG)

Continuous EEG was recorded from participants in the H-N group using a 64-channel BioSemi ActiveTwo system (BioSemi B.V., Amsterdam, The Netherlands). Ag/AgCl active electrodes with integrated preamplifiers were mounted on an elastic cap according to the international 10–10 system. Six participants were excluded because technical problems during acquisition resulted in unusable recordings, leaving a final sample of 31 participants for all EEG analyses. In the BioSemi ActiveTwo system, the conventional ground electrode is replaced by a Common Mode Sense (CMS) active electrode and a Driven Right Leg (DRL) passive electrode, which together form a feedback loop that stabilises the common-mode voltage. EEG signals were digitised at 1024 Hz with 24-bit resolution using ActiView software (BioSemi B.V.). Horizontal and vertical electrooculographic (EOG) activity was simultaneously recorded using four additional Ag/AgCl electrodes placed at the outer canthi of both eyes and above and below the left eye.

EEG preprocessing was performed in BrainVision Analyzer 2 (Brain Products, Gilching, Germany). Continuous data were downsampled to 200 Hz, band-pass filtered between 0.5 and 90 Hz, and notch filtered at 50 Hz to attenuate power-line interference. Noisy channels were identified by visual inspection and interpolated using spherical spline interpolation (spline order = 4). The data were subsequently re-referenced to the common average. Ocular artefacts, such as those associated with blinks and saccades, were identified and removed using independent component analysis (ICA). The cleaned EEG recordings were exported to Brainstorm (Tadel et al., 2011) for source-space analysis and segmented into epochs time-locked to the auditory cue signalling the onset of shape exploration. Each epoch comprised a 7-s pre-exploration static period followed by a 30-s active tracing period (Fig. 2C). All epochs and channels were visually inspected before analysis. Epochs containing non-physiological artefacts (e.g., high-amplitude transients, excessive movement artefacts, or widespread channel contamination) and persistently noisy channels were rejected. After these rejections, an average of 17.4 ± 2.8 (mean ± SD) artefact-free epochs per participant remained for analysis.

Cortical source reconstruction was performed using the Colin27 anatomical template (Montreal Neurological Institute) and the standard BioSemi 64-channel electrode layout. The forward model was computed using a three-shell spherical head model (scalp, outer skull and inner skull), which accounts for conductivity differences between tissue compartments (Hallez et al., 2007). The cortical source space comprised 15,002 vertices. Source activity was estimated using minimum norm estimation (MNE) with unconstrained dipole orientations and dynamic statistical parametric mapping (dSPM) noise normalisation. The three orthogonal dipole components at each vertex were retained for spectral analysis.

Spectral analysis was performed across the entire cortical surface using Welch’s method with 1-s Hamming windows and 50% overlap (Welch, 1967). Two analysis windows were defined for each trial. The resting window (4–9 s) was selected within the static pre-exploration period, and the active tracing window (12–38 s) was selected within the shape-exploration period. Both intervals were measured relative to trial onset and correspond to −6 to −1 s and 2 to 28 s relative to the tracing-onset cue, respectively. These windows were chosen to reduce the influence of transient auditory-evoked responses associated with the onset and offset cues while accommodating inter-individual variability in movement onset. Power spectral density (PSD) was estimated separately for the three orthogonal dipole components within each window. For each participant and period, the component PSDs were summed at each vertex and frequency and averaged across retained trials.

We focused on beta-band activity because of its established involvement in sensorimotor processing, with reductions in beta power often accompanying increased engagement of sensorimotor cortical regions (Cheyne et al., 2003; Pfurtscheller & Lopes da Silva, 1999). For each participant, PSD values were therefore averaged across beta-band frequency bins (15–29 Hz), separately for the resting and tracing periods. Group-level analyses then compared beta-band power between these periods across the cortical surface.

Finally, we examined whether task-related beta-power modulation during tracing was associated with subsequent performance in the reproduction and recognition tasks. Three regions of interest (ROIs) were defined a priori in each hemisphere based on the model of haptic object recognition proposed by Lacey and Sathian (2014). This model proposes that, during haptic perception of unfamiliar objects, the lateral occipital complex (LOC) is engaged through a predominantly bottom-up network involving the somatosensory cortex (S1) and the intraparietal sulcus (IPS). S1, located in the postcentral region, processes both cutaneous and proprioceptive information (Kim et al., 2015). The IPS is thought to contribute to the processing of the spatial relationships between object parts and their integration into a global shape representation, whereas the LOC is proposed to support object representations that are accessible across sensory modalities and independent of viewpoint (Lacey & Sathian, 2014). The S1 and LOC ROIs were defined using the Desikan–Killiany atlas (Desikan et al., 2006), whereas the IPS ROI, which is not present in the Desikan–Killiany atlas, was defined using the Destrieux atlas (Destrieux et al., 2010) (see Fig. 5). Following the whole-cortex cluster-based analysis, the anatomical overlap of these predefined ROIs with the significant beta-band clusters was quantified as ROI coverage, defined as the percentage of vertices comprising each ROI that fell within a significant cluster:

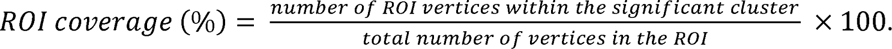

For each participant, mean beta-band power was extracted from each anatomical ROI, separately for the left and right hemispheres and for the tracing and resting periods. Task-related modulation relative to the resting baseline was calculated as 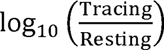. The resulting ROI-specific modulation measures were subsequently used to examine their relationship with shape reproduction and recognition accuracy.

### Statistical analysis

Statistical analyses were conducted using JASP (version 0.95.4) and RStudio (version 2025.05.1+513). Statistical significance was set at p < .05. Detailed statistical results are reported in the Supporting Information (Tables S1–S18).

Behavioural measures of tracing performance (i.e., the distance-to-segment index, number of shape traversals, mean tracing speed, mean normal force, and jerk index) were compared among the H-N, H-I, and VH-N groups using one-way ANOVA. Normality of the data was assessed using the Shapiro–Wilk test, and homogeneity of variance was assessed using Levene’s test. When the assumption of homogeneity of variance was met, classical one-way ANOVA was performed. Significant omnibus effects were followed by Tukey’s honestly significant difference (HSD) post hoc tests.

When homogeneity of variance was violated, Welch’s one-way ANOVA was used instead, followed by Games–Howell post hoc comparisons when significant omnibus effects were found. Effect sizes are reported as partial eta squared (η²p) for omnibus tests and Cohen’s d for pairwise comparisons.

Shape reproduction performance was quantified using the GFS score, which measures the similarity between the reproduced shape and the Reference shape. Differences among the five experimental groups (RC, H-N, H-I, VH-N, and V-N) were assessed using a one-way ANOVA followed by Tukey’s HSD post hoc comparisons.

Recognition performance was evaluated using two measures. First, the number of candidate shapes selected by each participant, used as a measure of recognition confidence (fewer selected shapes indicating greater confidence), was compared across groups using a Kruskal–Wallis test followed by Holm-corrected Dunn’s post hoc comparisons. Second, overall recognition performance was quantified using a GFS-based recognition score, calculated as the mean GFS similarity of the one to three shapes selected by each participant. Group differences were analysed using a Kruskal–Wallis test followed by Holm-corrected Dunn’s post hoc comparisons. Effect sizes for non-parametric analyses are reported as rank eta squared (rank η²) for omnibus tests and rank-biserial correlation (rrb) for pairwise comparisons.

For EEG source analyses, differences between resting and tracing periods were assessed using two-sided dependent-samples t-tests within a cluster-based permutation framework, implemented in FieldTrip (Oostenveld et al., 2011) through Brainstorm (Tadel et al., 2011). For each source point, the t-statistic was calculated across participants, and samples exceeding the cluster-forming threshold of p < .05 were grouped according to spatial adjacency. Cluster significance was determined using the maximum-sum (maxsum) statistic, with 5,000 random permutations used to generate the null distribution and control the family-wise error rate across the cortical source space. Clusters with corrected p < .05 were considered statistically significant.

To assess the relationship between ROI-specific beta-power modulation and subsequent memory performance, Spearman’s rank correlations (rho) were computed separately for the reproduction and recognition tasks. For each task, beta-power modulation during tracing relative to baseline was correlated with the corresponding GFS scores in each of the six a priori defined ROIs. P-values were adjusted using the Benjamini–Hochberg false discovery rate (FDR) procedure, applied separately to the six reproduction correlations and the six recognition correlations.

## Results

### Haptic exploration was shaped by memorisation instructions and visual feedback

Before assessing whether haptic information alone could support the formation of an internal representation of the traced shape, we first compared the kinematic and force characteristics of exploration in the H-N group with those of groups receiving memorisation instructions or visual feedback (Fig. 3). Welch’s ANOVA revealed a significant effect of Group on the distance-to-segment index (F_(2, 36.49)_ = 49.81, p < .001, η²p = .32; Fig. 3A). Games–Howell post hoc comparisons revealed no difference between the H-N and H-I groups (global mean = 1.43, p = .99), indicating that memorisation instructions did not affect tracing accuracy during purely haptic exploration. In contrast, the availability of visual feedback improved tracing accuracy, with the VH-N group exhibiting a significantly lower distance-to-segment index (mean = 1.13) than both H-N and H-I (both p < .001).

**Figure 3.**
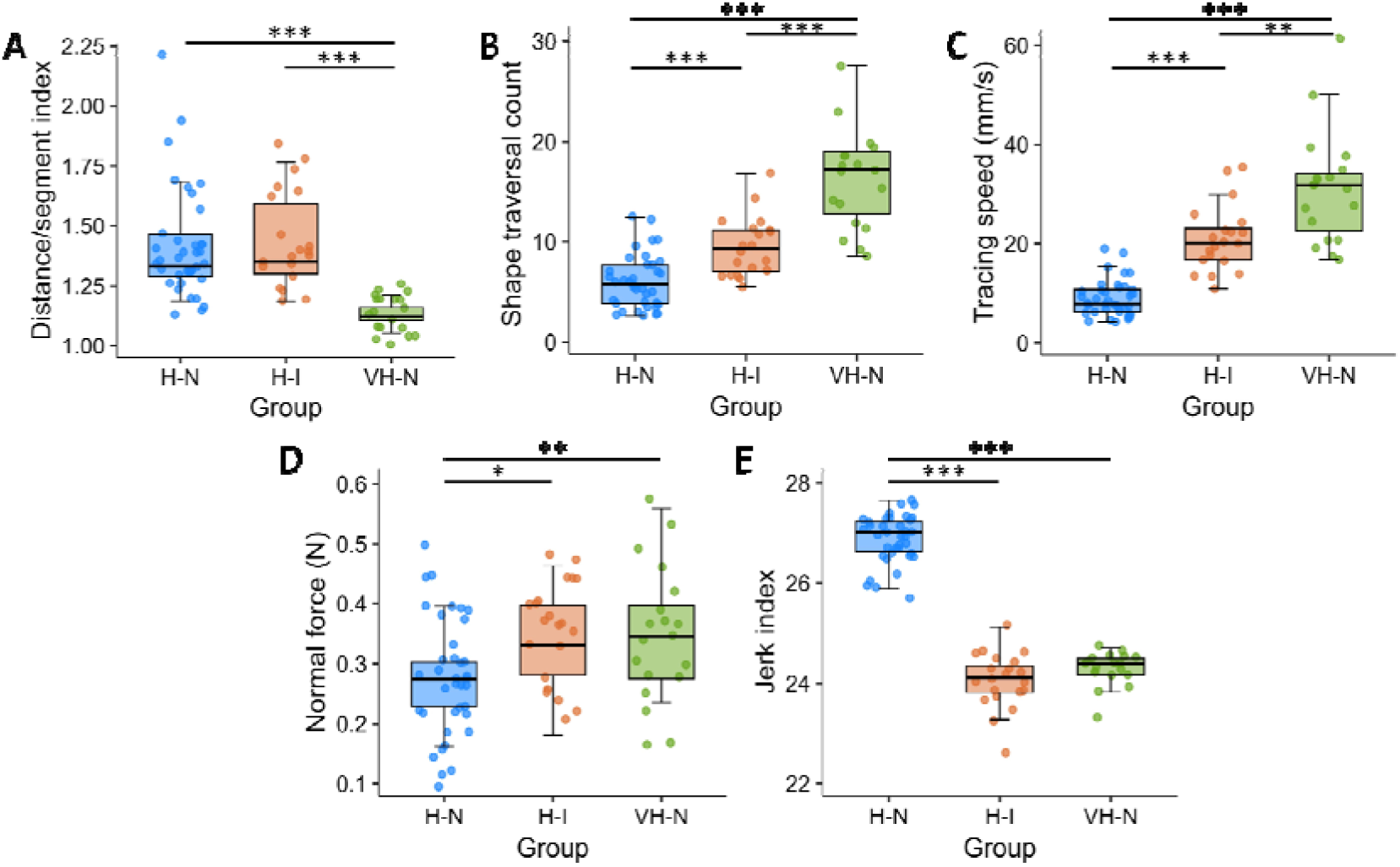
Kinematic and force characteristics of the exploratory phase. Boxplots showing the kinematic and force measures averaged across the tracing trials for the H-N (blue), H-I (orange), and VH-N (green) groups. Panels show **A** distance-to-segment index, **B** number of Reference shape traversals, **C** mean tracing speed (mm/s), **D** normal force (Fz) applied to the surface, and **E** jerk index. Central lines indicate medians, boxes the interquartile range (IQR), whiskers extend to 1.5 × IQR, and individual participants are shown as overlaid data points. * p < .05, ** p < .01, *** p < .001.

A one-way ANOVA revealed a significant effect of Group on tracing speed (F_(2,73)_ = 88.31, p < .001, η²p = .71; Fig. 3C). Tukey post hoc comparisons showed that the H-N group traced the shape significantly more slowly than both the H-I (p < .001, Cohen’s d = −2.43) and the VH-N (p < .001, Cohen’s d = −3.50) groups. The H-I group also traced significantly more slowly than the VH-N group (p = .004, Cohen’s d = −1.01). Consistent with these differences in tracing speed, the number of Reference shape traversals also differed significantly across groups (Welch’s ANOVA: F_(2, 33.55)_ = 31.18, p < .001, η²p = .57; Fig. 3B). Games–Howell post hoc comparisons showed that the H-N group completed fewer traversals (mean = 6.01) than both the H-I (mean = 9.49; p < .001) and VH-N (mean = 17.11; p < .001) groups, whereas the H-I group completed fewer traversals than the VH-N group (p < .001).

Forces exerted on the surface also differed between groups (Fig. 3D–E). One-way ANOVAs revealed significant effects of Group for both normal force (F_(2, 74)_ = 7.20, p = .001, η²p = .16; Fig. 3D) and jerk index (F_(2, 74)_ = 329.6, p < .001, η²p = .90; Fig. 3E). Tukey-corrected post hoc comparisons showed that the H-N group applied significantly lower normal force (mean = 0.27 N) than both the H-I (p = .02, Cohen’s d = −0.77) and the VH-N (global mean = 0.34 N; p = .003, Cohen’s d = −0.95) groups, whereas no difference was observed between H-I and VH-N groups (p = .84). Likewise, the H-N group exhibited markedly higher jerk values than both the H-I (p < .001, Cohen’s d = 6.10) and VH-N (p < .001, Cohen’s d = 5.56) groups, indicating less smooth modulation of shear force during exploration. The jerk index did not differ significantly between the H-I and VH-N groups (p = .21).

### Haptic exploration supports incidental spatial encoding

Group differences in shape reproduction performance, quantified using the GFS similarity scores, were assessed using a one-way ANOVA. Before analysis, one reproduction in the V-N group was identified as an outlier, as its GFS similarity score fell more than three standard deviations below the group mean (Stevens, 2002) and was excluded from the reproduction analysis (Supporting Information; Fig. S5, participant V-N_20). Figure 4A shows the reproductions with the highest and lowest GFS similarity scores in each group (see Figs S1–S5 for all individual drawings). A significant main effect of Group was observed (F_(4, 115)_ = 19.35, p < .001, η²p = .40; Fig. 4B). Tukey post hoc comparisons showed that the H-N group achieved significantly higher GFS scores than the RC group (p < .001), demonstrating that haptic exploration without explicit memorisation instructions was sufficient to yield reproductions that exceeded the baseline performance observed in the absence of target-specific information. GFS scores did not differ between the H-N and H-I groups (p = .71), suggesting that explicit memorisation instructions did not improve shape reproduction. Remarkably, the H-N and H-I groups did not differ from the VH-N group (p = .14 and p = .86, respectively), indicating that adding visual information to haptic input did not improve reproduction similarity. In contrast, the V-N group showed significantly higher GFS scores than the H-N group (p = .001; Cohen’s d = −1.13), reflecting better reproduction when the shape was visually available without haptic exploration. Finally, all four exploration groups achieved significantly higher GFS scores than the RC group (all p < .001, mean Cohen’s d = −1.89), confirming that their reproductions were not arbitrary.

**Figure 4.**
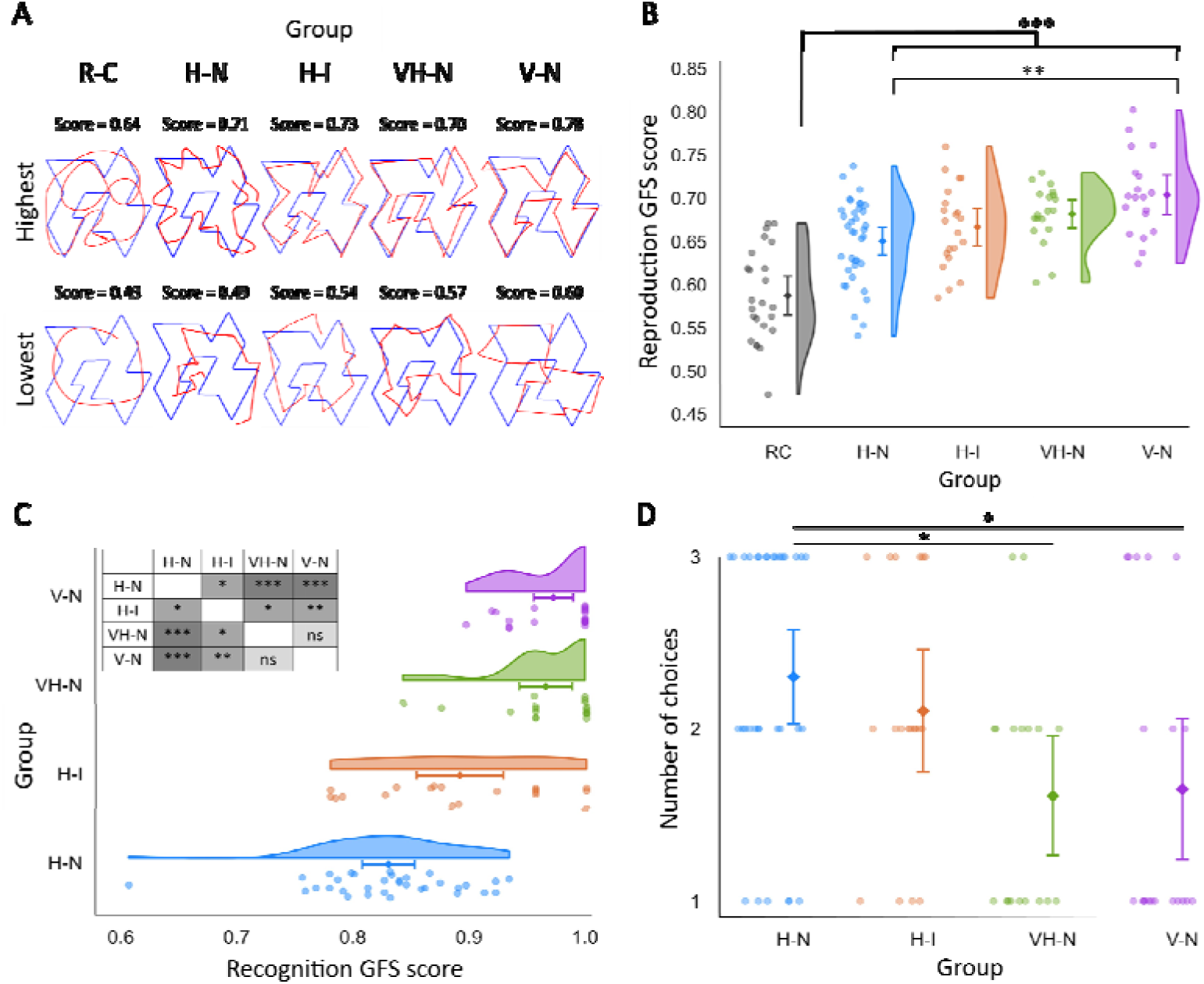
Shape reproduction and recognition performance across experimental groups. **A** Superimposition of the individual reproductions (red) with the highest (top row) and lowest (bottom row) GFS similarity scores within each group, overlaid on the Reference shape (blue). The corresponding GFS similarity score is indicated for each reproduction. **B** Raincloud plot showing the distributions of GFS similarity scores for drawings produced by the RC (grey), H-N (blue), H-I (orange), VH-N (green), and V-N (purple) groups. Higher scores indicate greater similarity between the drawings and the Reference shape. The GFS scores of the candidate shapes shown in Fig. 1C provide reference values for interpreting these similarities across groups. **C** Raincloud plot showing the GFS-based recognition score, calculated as the mean GFS similarity of the shapes selected by each participant. Higher scores indicate greater similarity of the selected shapes to the Reference shape (GFS = 1). The inset table summarises the post hoc comparisons. **D** Number of candidate shapes selected during the recognition task. Participants could select up to three shapes. In panels B, C and D, individual observations are shown as scatter points, with group means and their 95% confidence intervals overlaid. Panels B and C additionally display the half-violin density representing the distribution of the data. * p < .05, ** p < .01, *** p < .001; ns, not significant.

Recognition performance was quantified using a GFS-based recognition score, calculated as the mean similarity score of the one to three shapes selected by each participant. The GFS similarity scores of the candidate shapes shown in Fig. 1C provide a basis for interpreting these recognition scores reported below. A Kruskal–Wallis test revealed a significant effect of Group (χ^2^_(3)_ = 50.45, p < .001, rank η² = .55; Fig. 4C). Dunn’s post hoc comparisons showed that the H-N group (mean = 0.83) achieved significantly lower recognition scores than the H-I (mean = 0.86; p = .036; rrb = 0.42), VH-N (mean = 0.97; p < .001; rrb = 0.93), and V-N (mean = 0.97; p < .001; rrb = 0.97) groups. The H-I group also scored lower than both the VH-N (p = .011; rrb = 0.57) and V-N (p = .005; rrb = 0.62) groups, whereas the VH-N and V-N groups did not differ (p = .82). Thus, although the H-N group did not differ significantly from the other haptic groups in reproduction performance, it showed poorer recognition of the target shape. However, the mean GFS recognition scores of the H-N (0.83) and H-I (0.86) groups were substantially higher than the score of the least similar candidate shape (0.549) and approached the maximum value of 1.00, corresponding to the Reference shape.

The number of candidate shapes selected provided an index of recognition confidence, with fewer selections interpreted as greater confidence. A Kruskal–Wallis test revealed a significant effect of Group (χ^2^_(3)_ = 12.48, p = .006, rank η² = .11; Fig. 4D). Dunn’s post hoc comparisons revealed no significant difference between the H-N and H-I groups (mean = 2.20 for both groups; p = .82). In contrast, the H-N group selected significantly more candidate shapes than both the VH-N (mean = 1.61; p = .02; rrb = −0.47) and V-N (mean = 1.65; p = .02; rrb = −0.41) groups. No other pairwise differences were significant (all p ≥ .27). Thus, the lower recognition similarity in the H-N group was accompanied by the selection of more alternatives than in the VH-N and V-N groups, suggesting greater uncertainty in identifying the target shape.

### Beta-band power reduction during haptic exploration is associated with subsequent shape reproduction accuracy

Source-space spectral analyses were conducted in the H-N group to characterise cortical activity during haptic exploration in the absence of explicit memorisation instructions. Analyses focused on beta-band power (15–29 Hz), given the involvement of beta oscillations in sensorimotor processing (Pfurtscheller, 2001) and tactile working memory (Li Hegner et al., 2007). Beta power was quantified across the cortical surface and compared between the active tracing and the static pre-exploration resting phase (Fig. 5A). Tracing was associated with reduced beta power in two significant clusters, one in the left hemisphere (p = .0072; 3,162 vertices) and the other in the right hemisphere (p = .0124; 2,639 vertices). The left-hemisphere cluster showed a broad, predominantly fronto-temporo-parietal distribution, with additional posterior involvement. The right-hemisphere cluster showed a similarly widespread distribution, predominantly spanning fronto-parietal regions, with additional posterior involvement. To quantify the extent of overlap between the significant beta-band clusters and the atlas-defined ROIs selected based on the model proposed by Lacey and Sathian (2014), ROI coverage was calculated as the percentage of vertices within each ROI (S1, LOC, IPS) that fell within a significant cluster (table in Fig. 5A). All three ROIs overlapped the significant clusters bilaterally. Coverage was greater in the left than in the right hemisphere for S1 (65.71% vs. 46.13%), LOC (21.25% vs. 17.89%), and IPS (97.73% vs. 70.09%).

**Figure 5.**
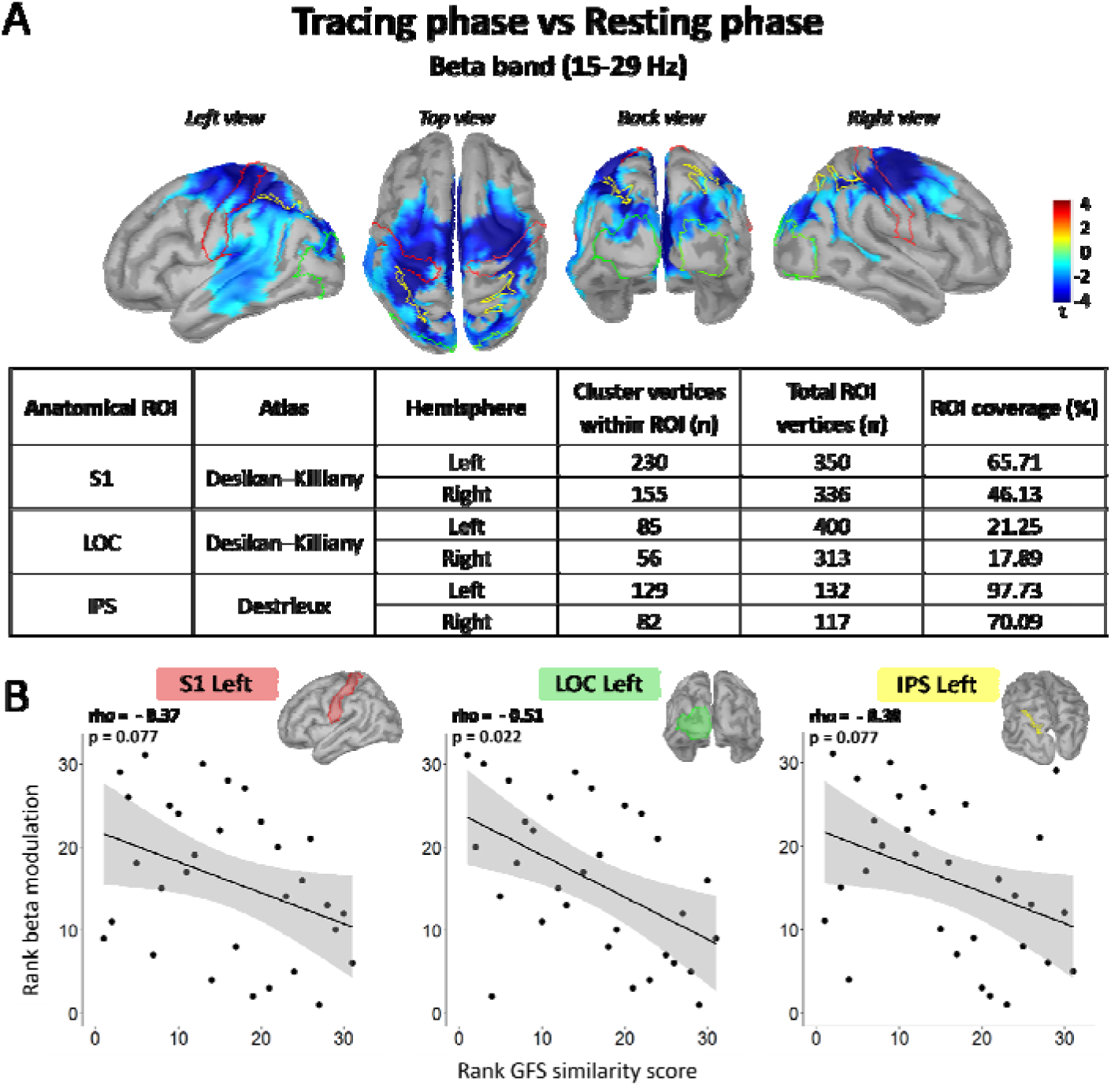
Cortical beta-band modulation during haptic exploration and its relationship with shape reproduction performance. **A** Source-space statistical maps showing significant clusters of reduced beta-band (15–29 Hz) power during tracing relative to the resting phase in the Haptic-Naïve (H-N) group. Differences were assessed using a two-sided cluster-based permutation test with a paired-samples t-statistic (5,000 permutations). Only clusters significant after cluster-level family-wise error correction (pFWE < .05) are displayed. The colour bar indicates t-values for the tracing-minus-resting contrast. Maps are shown on the MNI Colin27 cortical template (15,002 vertices), with 50% surface smoothing for visualisation, in left lateral, superior, posterior, and right lateral views. Coloured outlines indicate the predefined bilateral ROIs, S1 (red), LOC (green), and IPS (yellow). For each ROI and hemisphere, the table reports the anatomical atlas, the number of ROI vertices belonging to a significant cluster, the total ROI vertex count, and the corresponding ROI coverage (%). **B** Scatter plots showing ranked beta-power modulation during tracing relative to rest against ranked reproduction GFS similarity scores for the left S1, LOC, and IPS ROIs. Each point represents one participant. Solid lines show least-squares linear regression fits to the ranked data, and shaded areas represent 95% confidence intervals for the fitted mean. Spearman’s rank correlation coefficient (rho) and the corresponding p-value, adjusted for false discovery rate (FDR) across the six bilateral ROI-wise correlations, are reported for each plot. Negative coefficients indicate that greater beta-power reductions were associated with higher reproduction similarity scores. S1, primary somatosensory cortex; LOC, lateral occipital complex; IPS, intraparietal sulcus.

To examine whether beta-power modulation during exploration was associated with subsequent memory performance, we computed Spearman’s rank correlations between beta-power modulation during tracing relative to rest and GFS-based performance scores, separately for the reproduction and recognition tasks. For reproduction, FDR correction across the six ROI-wise correlations revealed a significant negative correlation in the left LOC (rho = −0.51, FDR-adjusted p = .022), indicating that greater beta-power reductions were associated with higher reproduction similarity scores (Fig. 5B). Negative correlations were also observed in the left S1 (rho = −0.37, uncorrected p = .038, FDR-adjusted p = .077) and left IPS (rho = −0.38, uncorrected p = .037, FDR-adjusted p = .077), but neither remained significant after correction for multiple comparisons. None of the right-hemisphere correlations reached statistical significance before correction (all uncorrected p ≥ .09).

For recognition, no significant correlations were observed between beta-power modulation and GFS-based recognition scores in any ROI, either before or after FDR correction applied separately across the six recognition correlations (Table S18).

## Discussion

The present study examined whether haptic exploration can support incidental encoding of the spatial structure of an unfamiliar shape without an explicit memorisation goal. By combining manual reproduction and visual recognition tasks with measures of exploratory behaviour, we assessed how memorisation instructions, visual feedback, and exploration mode influenced subsequent shape reproduction and recognition performance. We also characterised cortical activity during haptic exploration in the H-N group and examined its relationship with subsequent reproduction and recognition performance.

The most direct evidence for incidental spatial encoding comes from the comparison between the H-N and RC groups in the reproduction task. Participants in the RC group drew arbitrary shapes without prior exposure to the Reference, providing a baseline for reproduction in the absence of target-specific spatial information. Participants in the H-N group, who explored the Reference shape haptically without seeing it or anticipating the subsequent memory tasks, produced shapes that were significantly more similar to the Reference than those produced by the RC group. This finding indicates that haptic exploration without an explicit memorisation goal supported the retention of spatial information that could subsequently be used for manual reconstruction. Importantly, the H-I group, which was explicitly instructed to memorise the shape before haptic exploration, did not differ significantly from the H-N group in reproduction performance, indicating that no additional benefit of an explicit memorisation goal was detected for subsequent manual reconstruction. This similarity was observed despite the H-I group completing substantially more estimated contour traversals than the H-N group (9.49 vs. 6.01), raising the possibility that a spatial representation formed relatively early during contour tracing and remained available for subsequent reproduction without requiring deliberate memorisation or additional repetitions.

Although the explicit memorisation goal did not improve reproduction accuracy, it substantially altered how participants explored the shape. Compared with the H-N group, participants in the H-I group traced the contour faster and consequently completed more traversals, while also applying greater normal force. Faster tracing may reflect a strategy to reduce the interval between encounters with successive contour features. Greater normal force may likewise have increased mechanical stimulation of the skin, potentially enhancing the salience of tactile cues during exploration. Tracing accuracy did not differ significantly between the H-I and H-N groups, but the H-I group showed smoother modulation of tangential force magnitude, as indicated by lower jerk-index values. These differences in exploratory behaviour emerged despite identical contour-following instructions across groups, suggesting that the anticipated reproduction task influenced how participants performed their tracing movements. This finding is consistent with the goal-directed nature of haptic exploration, whereby exploratory movements can be adapted according to the information to be acquired (Lederman & Klatzky, 1987), perceptual constraints and stimulus characteristics (Katircilar & Drewing, 2023; Weiss & Flanders, 2011), and task goals (Sciutti et al., 2019). Specifically, Sciutti et al. (2019) showed that changing the goal of exploration from memorisation to comparison modified exploratory behaviour, even when the object and sensory modalities remained unchanged. Here, we further showed that even when the exploratory procedure is fixed, a prospective memory goal can modify how that procedure is executed, as reflected in tracing speed, normal force, and tangential force modulation. Notably, these changes in exploratory behaviour did not translate into more accurate reproduction, suggesting that the memorisation goal altered the manner in which tactile information was sampled rather than enhancing the resulting spatial representation.

Remarkably, providing direct visual feedback during haptic exploration also failed to improve subsequent reproduction relative to haptic-only exploration. Indeed, despite continuous visual access to the entire Reference shape and the exploring hand, reproduction performance in the VH-N group did not differ from that of either haptic-only group. Visual monitoring of the hand during exploration has previously been shown to impair haptic spatial perception, suggesting that viewing one’s own exploratory movements may capture attention and increase competition for limited working-memory resources, thereby interfering with the maintenance and rehearsal of haptic spatial information (Mueller et al., 2013). Unlike the free-exploration paradigm used by Mueller et al. (2013), the present study imposed a fixed exploratory procedure in which participants were instructed to follow the contour of the Reference shape with their index finger across successive trials. Like participants in the H-N group, those in the VH-N group were not informed about any subsequent tasks. It is therefore possible that, in the absence of an explicit memory goal, attentional resources during visuo-haptic exploration were primarily allocated to monitoring and controlling the ongoing tracing movement and to processing local contour features rather than the shape’s global configuration. Consistent with this interpretation, the VH-N group traced faster, completed more traversals, and followed the contour more accurately than the haptic-only groups, indicating that visual feedback facilitated online sensorimotor control without providing a corresponding advantage in subsequent memory-guided reproduction.

In contrast, the absence of visual feedback may have encouraged participants in the haptic-only groups to develop and use an internal representation of the shape to guide their exploratory movements. Reliance on this representation during exploration could have simultaneously supported online movement control and strengthened the representation available for subsequent reproduction. The comparable reproduction performance across the haptic-only and VH-N groups therefore suggests that more effective visually guided control did not necessarily confer an advantage in forming a representation for later memory-guided reproduction. On the other hand, the V-N group, which viewed the Reference shape without tracing it, reproduced it more accurately than the H-N group. This pattern raises the possibility that, even without an explicit memorisation goal, the absence of constraints associated with controlling a tracing movement allowed participants to allocate more attention to the Reference shape’s spatial configuration during visual exploration than during purely haptic exploration. The unfamiliar shape presumably offered limited affordances for manipulation or active interaction, which may have reduced the engagement of mechanisms involved in planning and controlling ongoing movements (Cisek & Kalaska, 2010; Gibson, 1966), likely reducing competition between movement control and the encoding of the shape’s spatial structure (Gunduz Can et al., 2017).

GFS-based shape recognition scores varied with both memorisation instructions and visual access to the shape. Although the H-N group scored lower than the other groups, its mean recognition score of 0.83 was close to the similarity scores of Shapes 4 and 5 (Fig. 1C), suggesting that incidental haptic exploration preserved substantial target-specific spatial information. Explicit memorisation instructions improved recognition performance, although no corresponding improvement in reproduction accuracy was detected. Recognition confidence nevertheless remained similarly low in the two haptic-only groups, which selected an average of 2.2 candidate shapes out of a maximum of three, compared with 1.63 in the visual groups. Although lower, this latter value was still relatively high, with some participants selecting two or even three candidates (see Fig. 4D), suggesting that even direct visual access to the shape did not guarantee confident recognition. Together, these findings highlight the potential importance of directed attention and explicit memorisation instructions for spatial encoding.

The distinct patterns of recognition and reproduction performance are consistent with the broader distinction between sensorimotor and cognitive modes of spatial encoding (Paillard, 1987). This distinction is supported by evidence that spatial information can guide action without being fully accessible to conscious perception (Blouin et al., 1995; Bridgeman et al., 1979; Goodale et al., 1991), whereas conscious perceptual access does not necessarily ensure its effective use for action (Goodale et al., 1994). Within this framework, our findings raise the possibility that explicit memorisation instructions favour the formation of cognitive representations of spatial information, improving subsequent recognition without necessarily enhancing the sensorimotor representations available for motor reproduction.

The formation of such cognitive representations may involve visual mediation, consistent with evidence that spatial information acquired without direct visual access to the target can engage visual and visuomotor representations (Darling et al., 2007; Manson et al., 2019), as well as with the image-mediation model of haptics (Lederman et al., 1990). The visual context available during haptic exploration may have contributed to this mediation. Although H-N and H-I participants could not see the Reference shape or their exploring hand, they could view the occluding shield and surrounding environment, which provided a stable visual reference frame. Such non-informative vision has been shown to facilitate haptic spatial perception (Newport et al., 2002) and may therefore have supported the organisation of spatial information acquired through touch.

The present findings should be interpreted in light of the distinct temporal characteristics of haptic and visual exploration. Whereas participants in the visual conditions had simultaneous access to the entire Reference shape, those in the haptic-only conditions acquired contour information progressively through encounters with local features. Haptic perception is inherently more sequential than visual perception (Picard & Monnier, 2009), and maintaining and integrating this information is thought to place substantial demands on working memory, particularly as stimulus complexity increases (Grunwald et al., 2001). In our experimental procedure, these demands were likely further increased by the sequential exploration, which consisted of 20 successive 30-s tracing periods. Despite this sequential acquisition, haptic tracing without an explicit memory goal provided sufficient spatial information to support subsequent reproduction and visual recognition. These findings are consistent with the possibility that information acquired through touch is maintained and progressively integrated through working-memory processes to construct a spatial representation (Moscatelli et al., 2014). Its accessibility and subsequent use may nevertheless depend on both task demands and the sensory information available during encoding, with explicit memorisation goals and visual information providing additional support for different aspects of memory performance.

Spectral analysis of source-reconstructed EEG characterised task-related beta-band modulation during haptic-only exploration in the H-N group, providing preliminary evidence of cortical activity that may contribute to incidental spatial encoding. Relative to the resting period, active tracing was associated with widespread beta-power reductions across bilateral frontoparietal and posterior regions, with additional temporal involvement in the left hemisphere. Such reductions are commonly observed during motor and somatosensory processing (Lebar et al., 2017; Pfurtscheller & Lopes da Silva, 1999) and have also been reported during tactile stimulation (Cheyne et al., 2003) and active touch (Park et al., 2019). Beta-power reductions have also been observed during stimulus presentation and retention in a tactile working-memory task (Li Hegner et al., 2007). Their broad cortical distribution is consistent with the widespread networks implicated in haptic perception (Sathian, 2016) and the maintenance of tactile spatial information in working memory (Schmidt & Blankenburg, 2018). In a tactile working-memory task using Braille-like patterns presented to the index finger, Schmidt and Blankenburg (2018) identified stimulus-specific representations in posterior parietal, premotor, and primary somatosensory cortices during a 12-s retention interval. At the same time, the present findings do not exclude a contribution of mental imagery during haptic exploration, even in the absence of an explicit memorisation goal. A recent activation likelihood estimation meta-analysis identified a distributed frontoparietal network associated with mental imagery, with partly distinct neural patterns for object and spatial imagery (Hawes et al., 2026). The posterior and parietal involvement observed here is therefore consistent with, but does not establish, a contribution from imagery-related processes.

The significant beta-band clusters overlapped the predefined bilateral S1, IPS, and LOC regions, which form part of the predominantly bottom-up network proposed for haptic perception of unfamiliar objects (Lacey et al., 2010; Lacey & Sathian, 2014). Within this network, the IPS may contribute to integrating spatial relationships between object parts, whereas the LOC may support a modality-independent representation of object shape. Source-space EEG studies have also identified feedforward beta-band interactions linking somatosensory, parietal, and occipital regions during tactile discrimination, with stronger coupling along certain pathways associated with greater perceptual accuracy (Adhikari et al., 2014). Although connectivity and directionality were not assessed here, these converging findings suggest that beta-band dynamics may contribute to the distributed processing of tactile information during active exploration. Bilateral LOC involvement is particularly relevant because this region has repeatedly been implicated in haptic processing without concurrent visual exploration (James et al., 2002). Its involvement here raises the possibility that occipital regions contributed to processing spatial structure acquired through touch. However, whether this involvement reflects visual mediation (Lederman et al., 1990) or the engagement of amodal spatial representations (Kennedy, 1993) remains unclear, as neither visual imagery nor interregional connectivity was directly assessed.

The relationship between beta-band modulation and subsequent reproduction performance further linked cortical activity during haptic exploration to the later use of the acquired spatial information. Greater reductions in beta-band power in the left LOC during tracing were associated with more accurate subsequent reproduction. Negative associations were also observed in the left S1 and IPS, although these did not survive FDR correction for multiple comparisons. No significant associations with reproduction performance were found in the right-hemisphere ROIs, and no significant correlations with recognition performance were observed in any ROI. These findings suggest that beta-band modulation during haptic exploration may reflect processes supporting the retention of spatial information for subsequent behaviour, consistent with its involvement in tactile working memory (Li Hegner et al., 2007). Given the established role of beta-band activity in sensorimotor processing (Cheyne et al., 2003; Pfurtscheller & Lopes da Silva, 1999), its relationship with subsequent memory performance may be particularly evident when retained information is used for manual reconstruction. Recognition performance may additionally depend on other neural modulations, offering one possible explanation for the absence of significant correlations with the beta-power changes examined here.

Taken together, the present findings suggest that active touch can leave a spatial trace that persists beyond exploration and subsequently influences behaviour and perception, even without an explicit memorisation goal. These findings build on the established contribution of touch to spatial perception (K. Johnson, 2001) and the guidance of ongoing movements (Moscatelli et al., 2019; Vlachou et al., 2025; Vlachou, Lafaverges, et al., 2026). The extent to which this spatial information can subsequently be accessed and used appears to depend, as our results suggest, on the conditions under which it is acquired, including the goals of exploration and the sensory information available. At the neural level, haptic-only exploration was associated with widespread reductions in beta-band power across a distributed somatosensory–parietal–occipital network, with modulation in the left LOC associated with subsequent reproduction performance. However, these beta-band changes cannot be attributed specifically to incidental spatial encoding, as EEG was recorded only in the H-N group, precluding a direct comparison with intentional encoding. The mechanisms underlying the behavioural and neural findings also remain uncertain, as neither attention nor encoding strategies were directly assessed or experimentally manipulated. Future studies combining neurophysiological recordings across encoding conditions with direct measures or manipulations of attention and encoding strategies could clarify how haptically acquired spatial information is encoded, maintained, and subsequently used to guide behaviour.

## Data availability statement

Detailed statistical outputs are provided in the Supporting Information. The datasets generated and analysed during the current study are available from the corresponding author upon reasonable request.

## Competing interests

The authors declare no competing interests.

## Author contributions

B.J., V.M-E., and M.L. conceived and designed the study. V.M-E. and K.S. collected and analysed the data. V.M-E. drafted the manuscript. V.M-E., B.J., M.L. reviewed and edited the manuscript. All authors approved the final version of the manuscript and agree to be accountable for all aspects of the work, ensuring that questions related to the accuracy or integrity of any part of the work are appropriately investigated and resolved. All persons designated as authors qualify for authorship, and all those who qualify for authorship are listed.

## Funding

The project leading to this publication has received funding from the French government under the “France 2030” investment plan managed by the French National Research Agency (reference: ANR-16-CONV000X / ANR-17-EURE-0029), from Excellence Initiative of AixMarseille University - A*MIDEX (AMX-19-IET-004), and the project COMTACT (ANR 2020-CE28-0010-03) and EMOTACT (ANR-24-CE28-0649) funded by the French “Agence Nationale de la Recherche” (ANR).

## Acknowledgements

We thank Dany Paleressompoulle and Eric Bergeret for fabricating and 3D-printing the surfaces, Franck Buloup for developing the Docometre software used for force data acquisition, and all participants for taking part in the study.

## Notes

### Competing Interest Statement

The authors have declared no competing interest.

