## Supporting Information for "Mapping the invisible: Incidental spatial encoding through haptic exploration"

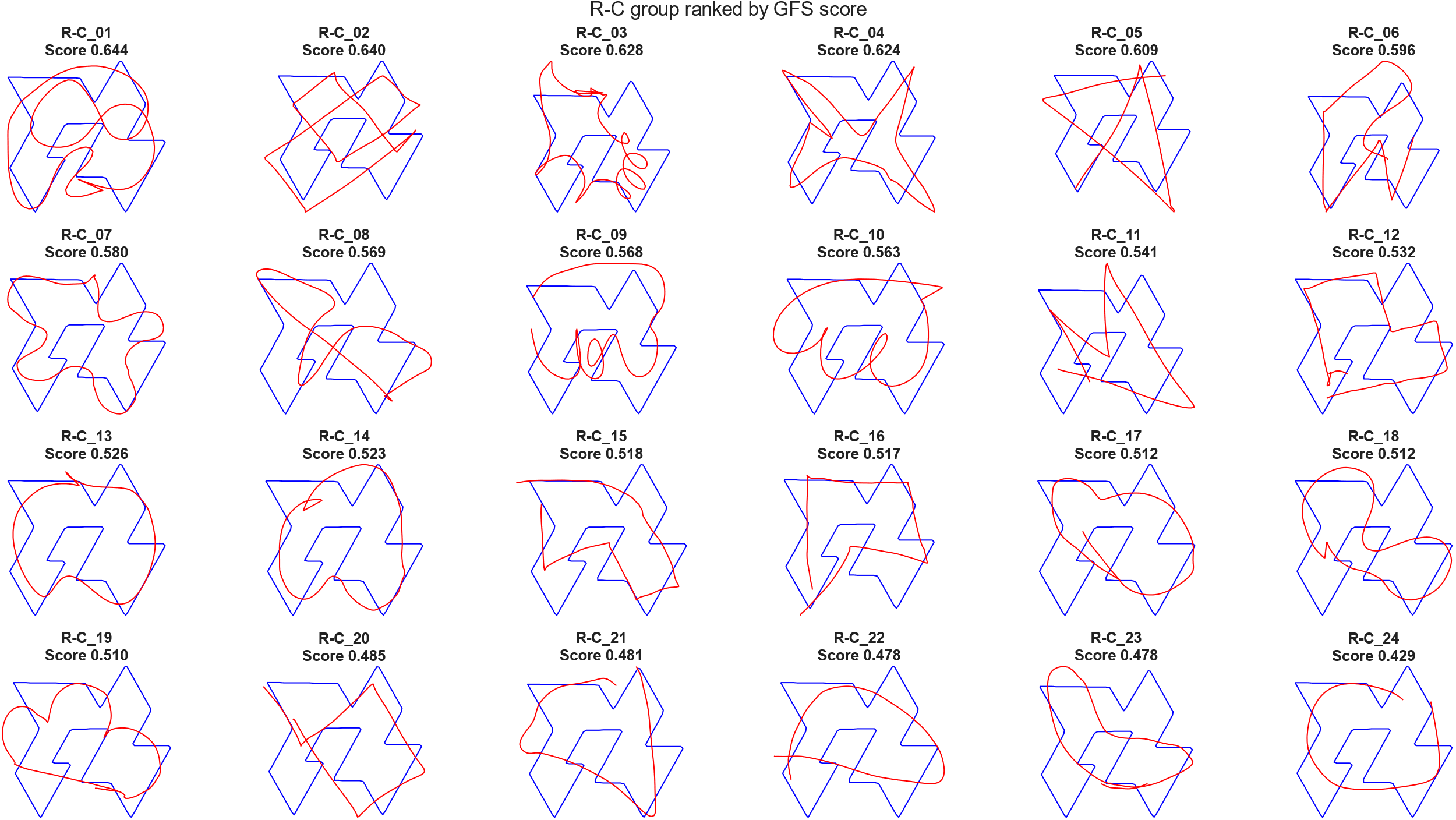

**Figure S1.** **Individual drawings in the Random-Control group.** Participants in the Random-Control (RC) group (n = 24) drew arbitrary closed shapes without prior exposure to the Reference shape. Individual drawings (red) are overlaid on the Reference shape (blue) and arranged in descending order of Geometric–Feature Similarity (GFS) score, from left to right and top to bottom. Participant labels and corresponding GFS scores are shown above each panel. Higher scores indicate greater similarity to the Reference shape.

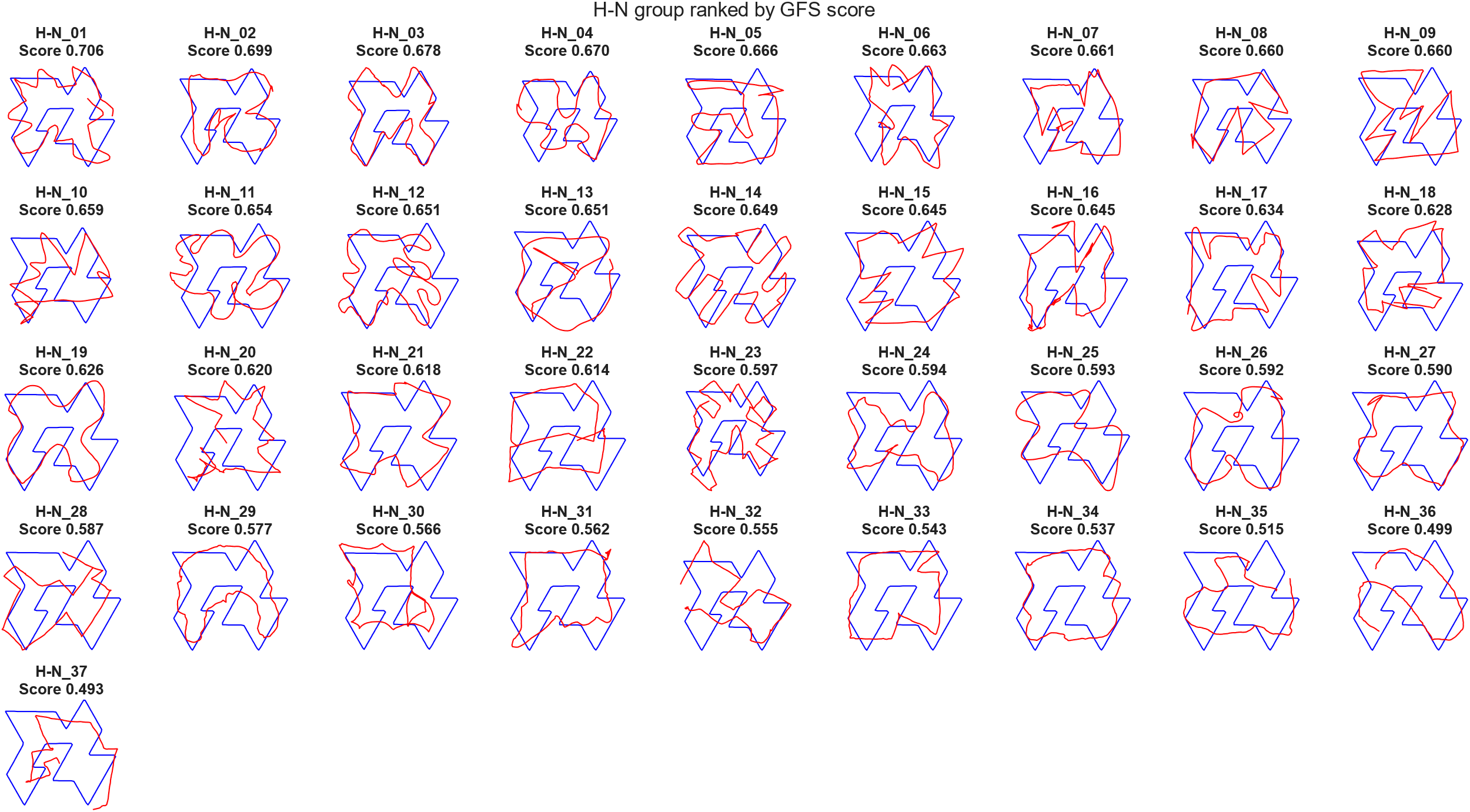

**Figure S2. Individual reproductions in the Haptic-Naïve group.** Participants in the Haptic-Naïve (H-N) group (n = 37) traced the Reference contour while it remained hidden from view, without advance knowledge of the subsequent memory tasks. Individual reproductions (red) are overlaid on the Reference shape (blue) and arranged in descending order of Geometric–Feature Similarity (GFS) score, from left to right and top to bottom. Participant labels and corresponding GFS scores are shown above each panel. Higher scores indicate greater similarity to the Reference shape.

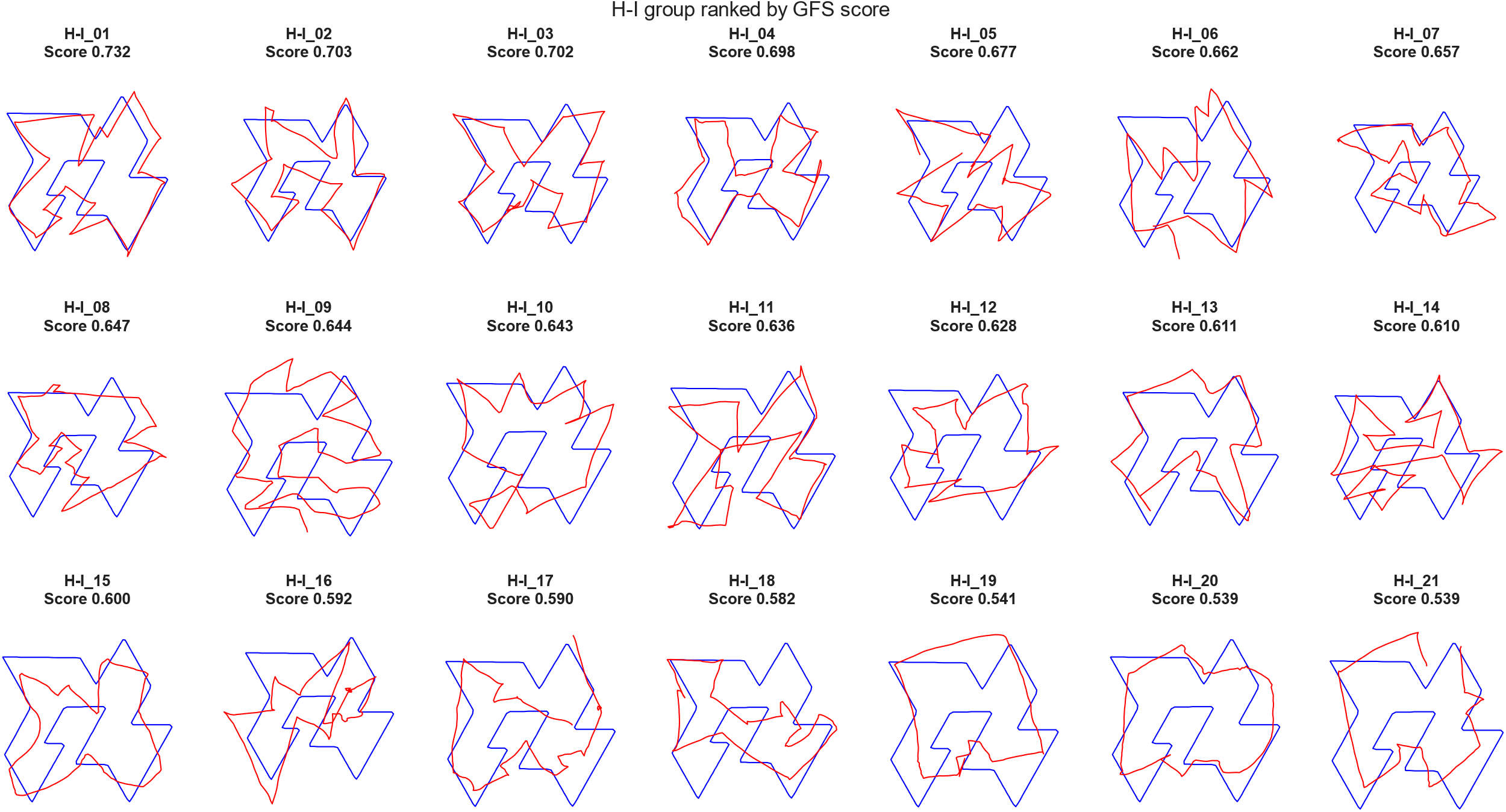

**Figure S3.** **Individual reproductions in the Haptic-Informed group.** Participants in the Haptic-Informed (H-I) group (n = 21) traced the Reference contour while it remained hidden from view, after receiving explicit instructions to memorise the shape. Individual reproductions (red) are overlaid on the Reference shape (blue) and arranged in descending order of Geometric–Feature Similarity (GFS) score, from left to right and top to bottom. Participant labels and corresponding GFS scores are shown above each panel. Higher scores indicate greater similarity to the Reference shape.

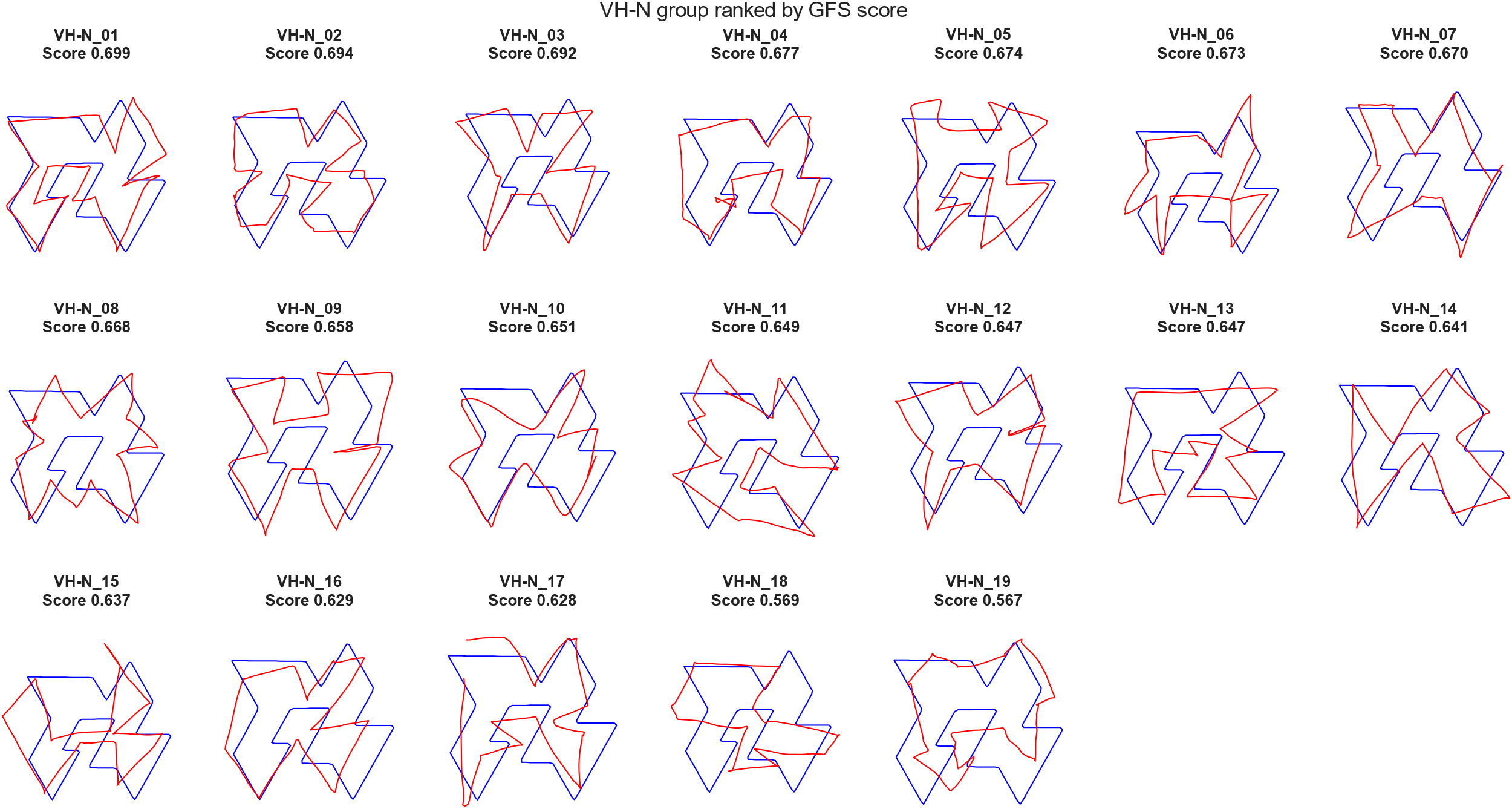

**Figure S4. Individual reproductions in the Visuo-Haptic-Naïve group.** Participants in the Visuo-Haptic-Naïve (VH-N) group (n = 19) traced the visible Reference contour without prior memorisation instructions. Individual reproductions (red) are overlaid on the Reference shape (blue) and arranged in descending order of Geometric–Feature Similarity (GFS) score, from left to right and top to bottom. Participant labels and corresponding GFS scores are shown above each panel. Higher scores indicate greater similarity to the Reference shape.

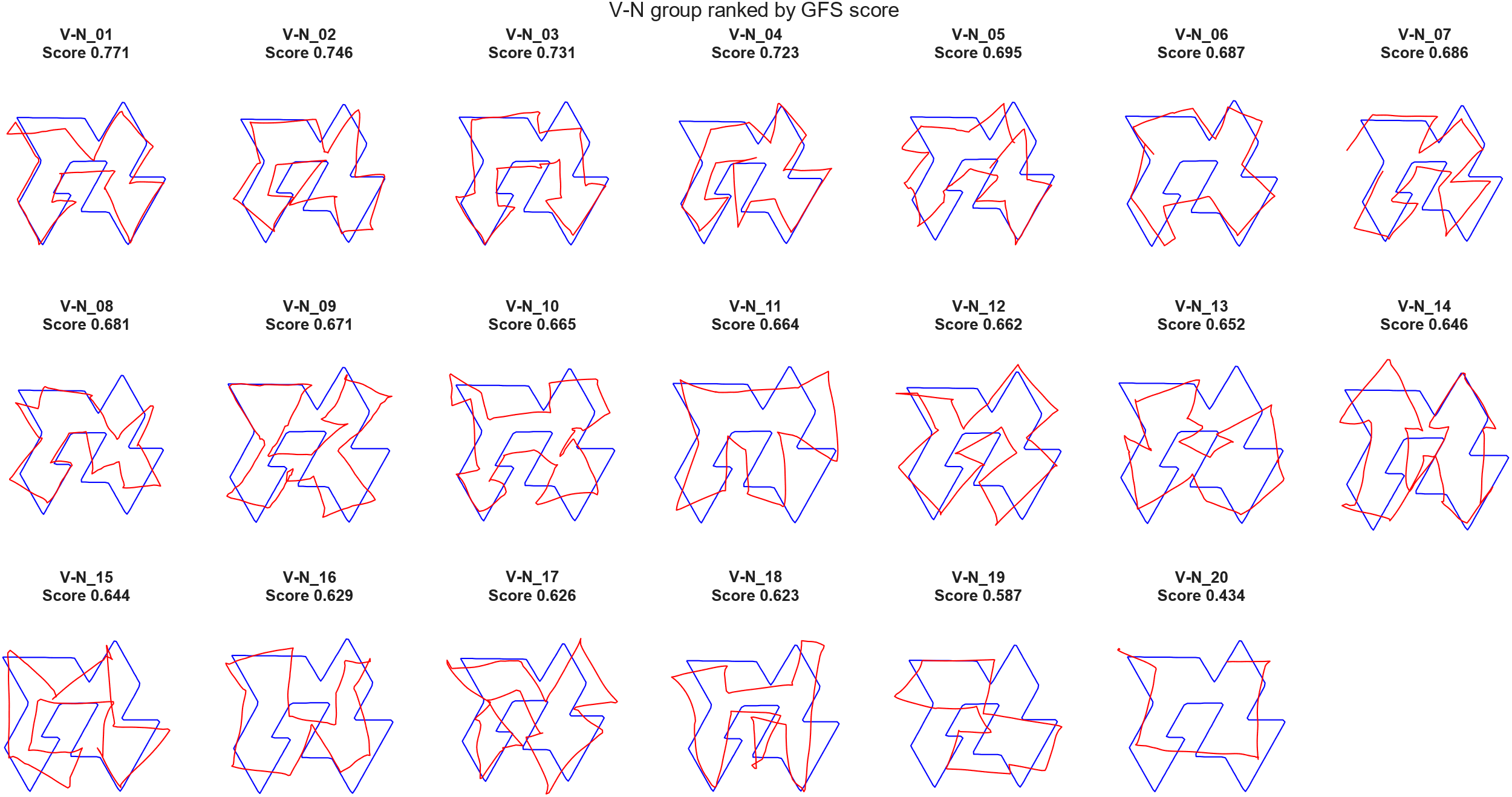

**Figure S5.** **Individual reproductions in the Visual-Naïve group.** Participants in the Visual-Naïve (V-N) group (n = 20) viewed the Reference shape without tracing it and without prior memorisation instructions. Individual reproductions (red) are overlaid on the Reference shape (blue) and arranged in descending order of Geometric–Feature Similarity (GFS) score, from left to right and top to bottom. Participant labels and corresponding GFS scores are shown above each panel. Higher scores indicate greater similarity to the Reference shape. The reproduction of participant V-N_20 was excluded from the statistical analysis of reproduction performance because its GFS score fell more than three standard deviations below the V-N group mean.

Table S1. **Distance/segment index:** ANOVA

| Homogeneity Correction | Cases | Sum of Squares | df | Mean Square | F | p | η²_p_ |
| --- | --- | --- | --- | --- | --- | --- | --- |
| Welch | Group | 1.25 | 2.00 | 0.63 | 49.81 | < .001*** | 0.32 |
|  | Residuals | 2.61 | 36.49 | 0.07 |  |  |  |

*** p < .001.

| Table S2. **Distance/segment index:** Games-Howell Post Hoc Comparisons - Group | | | | | |
| --- | --- | --- | --- | --- | --- |
| Comparison | Mean Difference | SE | t | df | p_tukey_ |
| H-N - H-I | -0.009 | 0.06 | -0.17 | 47.3 | 0.99 |
| H-N - VH-N | 0.29 | 0.04 | 7.45 | 40.29 | < .001*** |
| H-I - VH-N | 0.3 | 0.04 | 7.17 | 21.02 | < .001*** |
| *** p < .001. | | | | | |

**Table S1-S2.** Results of Welch’s one-way ANOVA and Games–Howell post hoc comparisons of the distance-to-segment index across the H-N, H-I, and VH-N groups. A significant effect of Group was observed. Pairwise comparisons revealed no significant difference between the H-N and H-I groups, whereas the VH-N group exhibited significantly lower index values than either haptic-only group.

| Table S3. **Number of shape traversals:** ANOVA | | | | | | | | |
| --- | --- | --- | --- | --- | --- | --- | --- | --- |
| Homogeneity Correction | Cases | Sum of Squares | | df | Mean Square | F | p | η²_p_ |
| Welch | Group | | 1548 | 2 | 773.9 | 31.18 | < .001*** | 0.57 |
|  | Residuals | | 1152 | 33.55 | 34.33 |  |  |  |
| *** p < .001.  *Note.*  Type III Sum of Squares | | | | | | | | |

| Table S4. **Number of shape traversals:** Games-Howell Post Hoc Comparisons - Group | | | | | |
| --- | --- | --- | --- | --- | --- |
| Comparison | Mean Difference | SE | t | df | p_tukey_ |
| H-N - H-I | -3.48 | 0.79 | -4.42 | 35.60 | < .001*** |
| H-N - VH-N | -11.10 | 1.53 | -7.25 | 21.19 | < .001*** |
| H-I - VH-N | -7.62 | 1.61 | -4.74 | 24.97 | < .001*** |
| *** p < .001. | | | | | |
| **Table S3-S4.** Results of Welch’s one-way ANOVA and Games–Howell post hoc comparisons of the estimated number of shape traversals across the H-N, H-I, and VH-N groups. A significant effect of Group was observed. Pairwise comparisons revealed fewer shape traversals in the H-N group than in both the H-I and VH-N groups, and fewer in the H-I group than in the VH-N group (all p < .001). | | | | | |

| Table S5. **Tracing speed:** ANOVA | | | | | | |
| --- | --- | --- | --- | --- | --- | --- |
| Cases | Sum of Squares | df | Mean Square | F | p | η²_p_ |
| Group | 4.27 | 2 | 2.13 | 88.31 | < .001*** | 0.71 |
| Residuals | 1.76 | 73 | 0.02 |  |  |  |
| *** p < .001. *Note.*  Type III Sum of Squares | | | | | | |

| Table S6. **Tracing speed:** Post Hoc Comparisons - Group | | | | | | | |
| --- | --- | --- | --- | --- | --- | --- | --- |
|  |  | Mean Difference | SE | df | t | Cohen's d | p_Tukey_ |
| (H-N) | (H-I) | -0.38 | 0.04 | 73 | -8.75 | -2.43 | < .001*** |
|  | (VH-N) | -0.54 | 0.04 | 73 | -12.41 | -3.5 | < .001*** |
| (H-I) | (VH-N) | -0.17 | 0.05 | 73 | -3.35 | -1.01 | .004** |
| ** p < .01, *** p < .001. | | | | | | | |
| *Note.*  P-value adjusted for comparing a family of 3 estimates. | | | | | | | |

**Table S5-S6.** Results of the one-way ANOVA and Tukey HSD post hoc comparisons of log-transformed tracing speed across the H-N, H-I, and VH-N groups. A significant effect of Group was observed. Pairwise comparisons revealed that the H-N group traced more slowly than both the H-I and VH-N groups (both p < .001), and that the H-I group traced more slowly than the VH-N group (p = .004).

| Table S7. **Normal force (Fz):** ANOVA | | | | | | |
| --- | --- | --- | --- | --- | --- | --- |
| Cases | Sum of Squares | df | Mean Square | F | p | η²_p_ |
| Group | 0.08 | 2 | 0.04 | 7.2 | .001** | 0.16 |
| Residuals | 0.40 | 74 | 0.005 |  |  |  |
| ** p < .01.  Note.  Type III Sum of Squares. | | | | | | |

| Table S8. **Normal force (Fz):** Post hoc comparisons - Group | | | | | | | |
| --- | --- | --- | --- | --- | --- | --- | --- |
|  |  | Mean Difference | SE | df | t | Cohen's d | p_Tukey_ |
| (H-N) | (H-I) | -0.06 | 0.02 | 74 | -2.8 | -0.77 | .02* |
|  | (VH-N) | -0.07 | 0.02 | 74 | -3.37 | -0.95 | .003** |
| (H-I) | (VH-N) | -0.01 | 0.02 | 74 | -0.57 | -0.18 | .84 |
| * p < .05, ** p < .01. | | | | | | | |
| *Note.*  P-value adjusted for comparing a family of 3 estimates. | | | | | | | |

**Table S7-S8.** Results of the one-way ANOVA and Tukey HSD post hoc comparisons of normal force (Fz) across the H-N, H-I, and VH-N groups. A significant effect of Group was observed. Pairwise comparisons revealed that the H-N group applied lower normal force than the H-I (p = .02) and VH-N (p = .003) groups, whereas no significant difference was found between the H-I and VH-N groups (p = .84).

| Table S9. **Jerk:** ANOVA | | | | | | |
| --- | --- | --- | --- | --- | --- | --- |
| Cases | Sum of Squares | df | Mean Square | F | p | η²_p_ |
| Group | 141.7 | 2 | 70.86 | 329.6 | < .001*** | 0.9 |
| Residuals | 15.91 | 74 | 0.22 |  |  |  |
| *** p < .001. *Note.*  Type III Sum of Squares | | | | | | |

| Table S10. **Jerk:** Post hoc comparisons - Group | | | | | | | |
| --- | --- | --- | --- | --- | --- | --- | --- |
|  |  | Mean Difference | SE | df | t | Cohen's d | p_Tukey_ |
| (H-N) | (H-I) | 2.83 | 0.13 | 74 | 22.32 | 6.1 | < .001*** |
|  | (VH-N) | 2.58 | 0.13 | 74 | 19.71 | 5.56 | < .001*** |
| (H-I) | (VH-N) | -0.25 | 0.15 | 74 | -1.70 | -0.54 | 0.21 |
| *** p < .001. | | | | | | | |
| *Note.*  P-value adjusted for comparing a family of 3 estimates. | | | | | | | |

**Table S9-S10.** Results of the one-way ANOVA and Tukey HSD post hoc comparisons of the jerk index across the H-N, H-I, and VH-N groups. A significant effect of Group was observed. Pairwise comparisons revealed higher jerk-index values in the H-N group than in both the H-I and VH-N groups (both p < .001), whereas no significant difference was found between the H-I and VH-N groups (p = .21).

| Table S11. **GFS similarity score:** ANOVA | | | | | | |
| --- | --- | --- | --- | --- | --- | --- |
| Cases | Sum of Squares | df | Mean Square | F | p | η²_p_ |
| Group | 0.17 | 4 | 0.04 | 19.35 | < .001*** | 0.4 |
| Residuals | 0.26 | 115 | 0.002 |  |  |  |
| *** p < .001. *Note.*  Type III Sum of Squares | | | | | | |

| Table S12. **GFS similarity score:** Post Hoc Comparisons - Group | | | | | | | |
| --- | --- | --- | --- | --- | --- | --- | --- |
|  |  | Mean Difference | SE | df | t | Cohen's d | p_Tukey_ |
| RC | (H-N) | -0.06 | 0.01 | 115 | -5.12 | -1.34 | < .001*** |
|  | (H-I) | -0.08 | 0.01 | 115 | -5.64 | -1.69 | < .001*** |
|  | (VH-N) | -0.09 | 0.01 | 115 | -6.52 | -2 | < .001*** |
|  | (V-N) | -0.12 | 0.01 | 115 | -8.04 | -2.47 | < .001*** |
| (H-N) | (H-I) | -0.02 | 0.01 | 115 | -1.27 | -0.35 | 0.71 |
|  | (VH-N) | -0.03 | 0.01 | 115 | -2.34 | -0.66 | 0.14 |
|  | (V-N) | -0.05 | 0.01 | 115 | -4 | -1.13 | .001** |
| (H-I) | (VH-N) | -0.01 | 0.01 | 115 | -0.99 | -0.31 | 0.86 |
|  | (V-N) | -0.04 | 0.01 | 115 | -2.47 | -0.78 | 0.1 |
| (VH-N) | (V-N) | -0.02 | 0.02 | 115 | -1.44 | -0.47 | 0.6 |
| ** p < .01, *** p < .001. | | | | | | | |
| Note.  P-value adjusted for comparing a family of 5 estimates. | | | | | | | |

**Table S11-S12.** Results of the one-way ANOVA and Tukey HSD post hoc comparisons of reproduction GFS similarity scores across the RC, H-N, H-I, VH-N, and V-N groups. A significant effect of Group was observed. Pairwise comparisons revealed higher scores in all four exploration groups than in the RC group (all p < .001), and higher scores in the V-N group than in the H-N group (p = .001). No other pairwise differences were significant (all p ≥ .10).

| Table S13. **Number of choices:** Kruskal-Wallis Test | | | | | | |
| --- | --- | --- | --- | --- | --- | --- |
|  | | | | | 95% CI for Rank η² | |
| Factor | Statistic | df | p | Rank η² | Lower | Upper |
| Group | 12.48 | 3 | 0.006** | 0.11 | 0.02 | 0.31 |

** p < .01.

| Table S14. **Number of choices:** Dunn's Post Hoc Comparisons - Group | | | | | |
| --- | --- | --- | --- | --- | --- |
| Comparison | z | W_i_ | W_j_ | r_rb_ | p_Holm_ |
| H-N - H-I | 0.83 | 55.26 | 49.39 | -0.15 | 0.82 |
| H-N - VH-N | 2.87 | 55.26 | 34.53 | -0.47 | .02* |
| H-N - V-N | 2.82 | 55.26 | 35.58 | -0.41 | .02* |
| H-I - VH-N | 1.84 | 49.39 | 34.53 | -0.36 | 0.27 |
| H-I - V-N | 1.75 | 49.39 | 35.58 | -0.32 | 0.27 |
| VH-N - V-N | -0.13 | 34.53 | 35.58 | -0.02 | 0.9 |
| * p < .05, ** p < .01. | | | | | |
| *Note.*  Rank-biserial correlation based on individual Mann-Whitney tests. | | | | | |

**Table S13-S14.** Results of the Kruskal–Wallis test and Holm-corrected Dunn post hoc comparisons of the number of candidate shapes selected during recognition across the H-N, H-I, VH-N, and V-N groups. A significant effect of Group was observed. Pairwise comparisons revealed no significant difference between the H-N and H-I groups (p = .82), whereas the H-N group selected more candidate shapes than both the VH-N and V-N groups (both p = .02). No other pairwise differences were significant (all p ≥ .27).

| Table S15. **Recognition score:** Kruskal-Wallis Test | | | | | | |
| --- | --- | --- | --- | --- | --- | --- |
|  | | | | | 95% CI for Rank η² | |
| Factor | Statistic | df | p | Rank η² | Lower | Upper |
| Group | 50.45 | 3 | < .001*** | 0.55 | 0.46 | 0.67 |

*** p < .001.

| Table S16. Recognition score**:** *Dunn's Post Hoc Comparisons - Group* | | | | | |
| --- | --- | --- | --- | --- | --- |
| Comparison | z | W_i_ | W_j_ | r_rb_ | p_Holm_ |
| H-N - H-I | -2.36 | 23.48 | 41.08 | 0.42 | .036* |
| H-N - VH-N | -5.59 | 23.48 | 65.83 | 0.93 | < .001*** |
| H-N - V-N | -6.04 | 23.48 | 67.73 | 0.97 | < .001*** |
| H-I - VH-N | -2.91 | 41.08 | 65.83 | 0.57 | .011* |
| H-I - V-N | -3.22 | 41.08 | 67.73 | 0.62 | .005** |
| VH-N - V-N | -0.23 | 65.83 | 67.73 | 0.04 | 0.82 |
| * p < .05, ** p < .01, *** p < .001. | | | | | |
| *Note.*  Rank-biserial correlation based on individual Mann-Whitney tests. | | | | | |

**Table S15-S16.** Results of the Kruskal–Wallis test and Holm-corrected Dunn post hoc comparisons of GFS-based recognition scores across the H-N, H-I, VH-N, and V-N groups. A significant effect of Group was observed. Pairwise comparisons revealed lower scores in the H-N group than in the H-I group (p = .036) and both visual groups (VH-N and V-N; both p < .001). The H-I group also scored lower than the VH-N (p = .011) and V-N (p = .005) groups, whereas no significant difference was found between the VH-N and V-N groups (p = .82).

Table S17. ROI correlations between beta-power modulation and reproduction similarity

| ROI | Hemisphere | n | rho | p | p_FDR_ |
| --- | --- | --- | --- | --- | --- |
| S1 | Left | 31 | -0.37 | .04* | .08 |
| S1 | Right | 31 | -0.31 | .09 | .13 |
| LOC | Left | 31 | -0.51 | .004** | .02* |
| LOC | Right | 31 | -0.24 | .2 | .2 |
| IPS | Left | 31 | -0.38 | .04* | .08 |
| IPS | Right | 31 | -0.27 | .14 | .17 |

* p < .05, ** p < .01.

**Table S17.** Spearman rank correlations between beta-band power during tracing relative to rest and subsequent reproduction GFS similarity scores in the H-N group. Negative coefficients indicate that larger beta-power reductions were associated with higher reproduction similarity. rho, Spearman’s rank correlation coefficient; n, number of participants; p, unadjusted p value; pFDR, p value adjusted using the Benjamini–Hochberg false discovery rate procedure across all six correlations. S1, primary somatosensory cortex; LOC, lateral occipital complex; IPS, intraparietal sulcus. Significance is based on pFDR < .05.

Table S18. ROI correlations between beta-power modulation and visual recognition performance

| ROI | Hemisphere | n | rho | p | p_FDR_ |
| --- | --- | --- | --- | --- | --- |
| S1 | Left | 27 | -0.31 | .054 | .32 |
| S1 | Right | 27 | -0.31 | .68 | .94 |
| LOC | Left | 27 | -0.51 | .78 | .94 |
| LOC | Right | 27 | -0.24 | 1 | 1 |
| IPS | Left | 27 | -0.38 | .46 | .91 |
| IPS | Right | 27 | -0.27 | .19 | .57 |

**Table S18.** Spearman’s rank correlations between beta-power modulation during tracing relative to the resting baseline and subsequent GFS-based recognition scores in the Haptic-Naïve (H-N) group. None of the six correlations was significant after FDR correction. rho, Spearman’s rank correlation coefficient; n, number of participants; p, unadjusted p value; pFDR, p value adjusted using the Benjamini–Hochberg false discovery rate procedure across all six correlations. S1, primary somatosensory cortex; LOC, lateral occipital complex; IPS, intraparietal sulcus. Significance is based on pFDR < .05.
